# A scalable, all-optical method for mapping synaptic connectivity with cell-type specificity

**DOI:** 10.1101/2025.06.25.661552

**Authors:** Maria V. Moya, William J. Cunningham, Jack P. Vincent, Tim Wang, Michael N. Economo

## Abstract

Single-cell transcriptomics has uncovered the vast heterogeneity of cell types that compose each region of the mammalian brain, but describing how such diverse types connect to form functional circuits has remained challenging. Current methods for measuring the presence and strength of synaptic connections principally rely on low-throughput whole-cell recording approaches. However, the development of optical tools for perturbing and observing neural activity—now notably including genetically encoded voltage indicators—presents an exciting opportunity to vastly increase the throughput of physiological connectivity mapping. Here, we combine massively parallel optical measurements of synaptic strength with a novel pipeline for thick-tissue spatial transcriptomics to assay synaptic connectivity motifs with high sensitivity, high throughput, and cell-type specificity. We apply this approach in the motor cortex to describe new cell-type-specific synaptic innervation patterns for long-range thalamic and contralateral input onto more than 1000 motor cortical neurons.

## Introduction

The function of a neural circuit is intimately related to its structure. Neuroanatomy has been instrumental for inspiring conceptual ideas and constraining computational models of circuit function in the mammalian cerebellum, hippocampus, basal ganglia, neocortex, and retina, among other areas^1–9^. The cell types comprising individual neural circuits have typically been identified and described in a painstaking fashion through a synthesis of information about their location, morphology, gene expression, electrophysiological characteristics, and patterns of activation *in vivo*. Consequently, identifying the connectivity motifs that govern how different cell types connect to each other and connect across distant regions has proven an enormously challenging endeavor. Recently, single-cell transcriptomic data have transformed our understanding of cell types, suggesting a much higher degree of cell type diversity than previously appreciated in most brain regions^10,11^. The rapid pace of progress in our understanding of neuronal cell types has also now far surpassed our ability to determine their local and long-range connectivity at a commensurate level of detail. Thus, effectively leveraging these advances into new insights about neural computation remains difficult.

This disconnect is due to a lack of scalable methods for determining the probability and strength of connections formed between transcriptomically defined cell types resident within neural circuits. “Paired-patch” electrophysiology has been used extensively and remains the “gold standard” for measuring local connectivity between pairs of neurons^12–14^. For identifying long-range connections, Channelrhodopsin-assisted circuit mapping (CRACM) combines optogenetic activation of axonal fibers with patch-clamp electrophysiology to read out postsynaptic responses^15,16^. These paradigms are labor intensive, difficult to scale, and thus cannot be readily used to determine connectivity in circuits containing many cell types. The rapid improvement in genetically encoded voltage indicators (GEVIs) offers an attractive alternative to patch-clamp electrophysiology in these paradigms^17–20^, opening the door for highly parallelized measurements of connectivity. However, this requires the detection of small, subthreshold synaptic potentials, which is challenging due to the limited signal-to-noise (SNR) of GEVI imaging.

Here, we describe an optical method for assaying synaptic connectivity that is both highly scalable and compatible with spatial transcriptomics for cell type identification. By combining optogenetics and GEVI imaging, we demonstrate that postsynaptic potentials (PSPs) can be optically elicited and detected in a sensitive and highly parallel manner across large populations of neurons, presenting a scalable alternative to patch-clamp electrophysiology for efficiently revealing circuit connectivity. We combine all-optical measurements of synaptic connectivity with a custom protocol for thick-tissue multiplexed fluorescence in situ hybridization (mFISH) to provide a Multimodal Optical Strategy for Assaying Identity and Connectomics (MOSAIX). We apply MOSAIX to map long-range thalamic and callosal inputs onto over 1000 transcriptomically identified cells in the motor cortex. Surprisingly, we find that long-range inputs are integrated with a high degree of specificity when cell types are precisely defined, such that even transcriptomically related cell types are often selectively targeted in a pathway-specific manner.

## Results

### Sensitive and scalable optical detection of synaptic potentials

GEVIs have principally been utilized for the detection of action potentials^21–25^. Another promising application is the detection of subthreshold PSPs, which would permit measuring synaptic connectivity in a highly parallel fashion^17,26,27^. However, the limited SNR of GEVI imaging can preclude the detection of PSPs, which are small by comparison. To determine the sensitivity with which PSPs could be detected using GEVI imaging, we expressed the blue-shifted opsin, CheRiff^28^, in cells of the motor thalamus (“MThal”) and the chemigenetic GEVI, Voltron2^18^, sparsely (2-5% of neurons) across the anterior-to-posterior extent of the primary motor cortex (**Figure 1a,b**). We performed simultaneous whole-cell electrophysiology and widefield voltage imaging of Voltron2-expressing (Voltron2+) neurons (**Figure S1a**) while optically activating CheRiff+ MThal axons to elicit synaptic potentials in the motor cortex *in vitro* (**Figure 1c,d**). Labeling Voltron2 with the JaneliaFluor 585 dye^29^ (Voltron2_JF585_) allowed spectrally separable voltage imaging and optogenetic activation.

**Figure 1.**
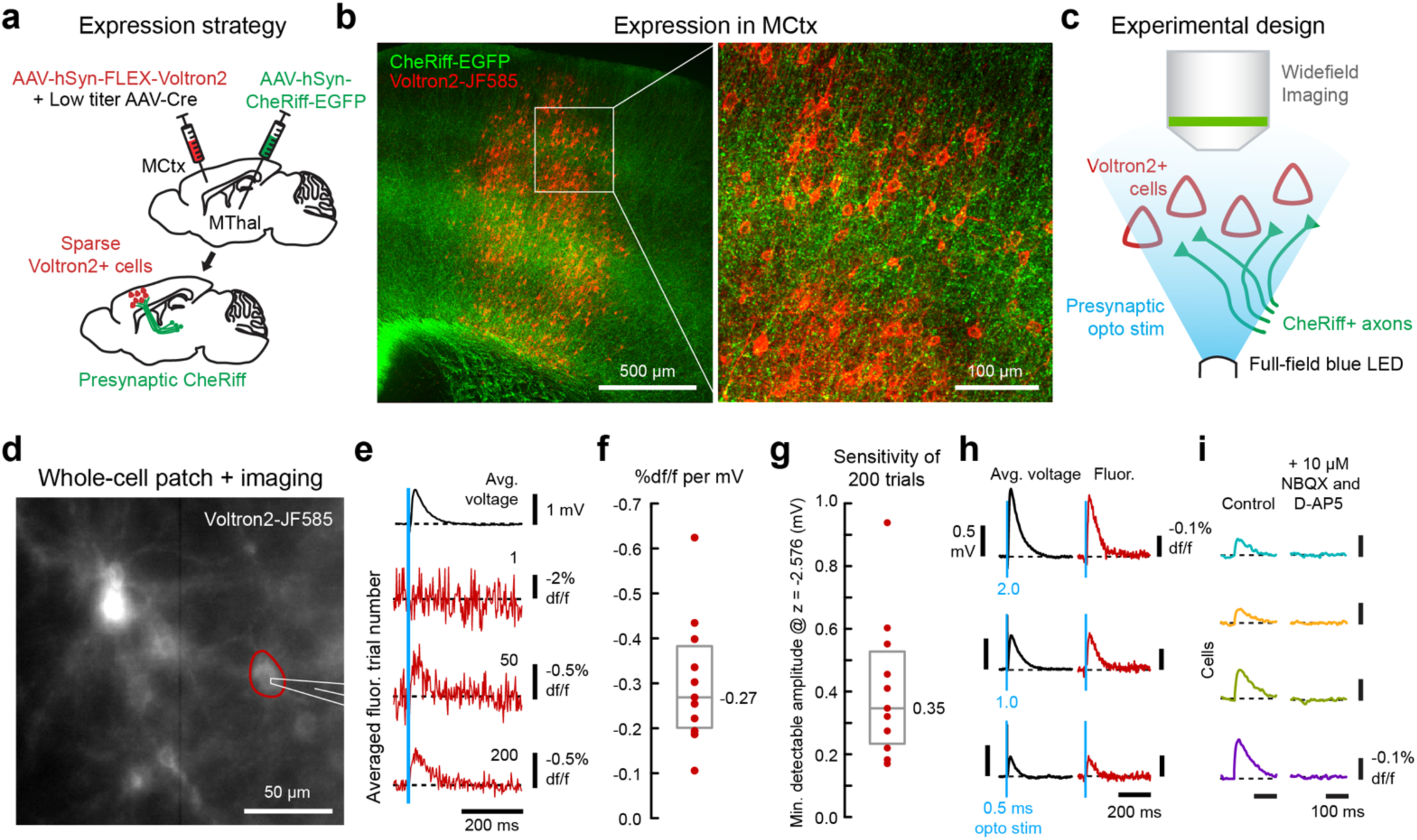
Sensitivity of voltage imaging for detection of postsynaptic potentials. (**a**) Schematic showing viral delivery and expression strategy for targeting CheRiff-EGFP to presynaptic neurons in the motor thalamus (MThal) and for sparsely expressing Voltron2 in postsynaptic neurons in the primary motor cortex. (***b***) Z-projected confocal image and inset of CheRiff-EGFP+ MThal axons (green) and Voltron2-JF585+ neurons (red) across 10 µm of motor cortex. (***c***) Schematic showing the experimental design for all-optical connectivity mapping. Acute slices containing CheRiff-positive MThal axons and Voltron2-expressing neurons are imaged on a widefield microscope. Presynaptic MThal axons are activated using a full-field blue LED. (***d***) Widefield image of Voltron2-JF585 signal in an acute slice, with a Voltron2+ neuron (red) patched for simultaneous whole-cell recording. White lines highlight location of patch pipette. (***e***) Postsynaptic potentials (PSPs) measured using whole-cell voltage recording (top, black) and fluorescent timeseries extracted from cell in ***d*** (red outline and traces). Averaging PSP profiles across repetitive photoactivation trials (1, 50, and 200 trial-averages are shown) increases the response signal-to-noise. (***f***) Box and whisker plot showing the percent change in fluorescence observed for Voltron2-JF585 for a 1 mV change in membrane voltage across all cells patched (red dots; n = 11; median = –0.27%, IQR –0.38 to – 0.20%). (***g***) Box and whisker plot showing the minimum PSP amplitude detectable (with statistical confidence, z ≤ –2.576) over noise when averaging 200 photoactivation trials across all cells patched (red dots; n = 11; median = 0.35 mV, IQR 0.23 to 0.53 mV). (***h***) Whole-cell (black) and fluorescent (red) traces from one patched neuron showing that PSP amplitude can be modulated by employing differing stim pulse durations during all-optical connectivity mapping. (***i***) Trial-averaged traces from 4 neurons in the same field-of-view, showing that responses are abolished in the presence of glutamatergic blockers (10 µM NBQX and D-AP5), confirming their synaptic origin.

Photoactivation of populations of afferent axons can typically evoke PSPs ranging from tens of millivolts down to single-millivolt amplitudes in connected cortical neurons^30–32^. GEVI measurements of PSPs evoked using brief blue light stimulation pulses were noisy, such that millivolt-scale PSPs could not be reliably detected on single trials (**Figure 1e**). However, averaging optical measurements across repeated photostimuli reduced noise, revealing PSPs with high SNR (**Figure 1e**). To calibrate optical voltage measurements, we characterized the relationship between fluorescence and membrane voltage by measuring voltage simultaneously with GEVI imaging and patch clamp electrophysiology. For all cells, this relationship was linear with a median slope of –0.27%/mV (**Figure 1f**; IQR –0.38 to –0.20%; n = 11), as expected for Voltron2, a negative-going GEVI ^18^. Based on the F-V relationship and noise level measured in each cell, we calculated the minimum PSP amplitude detectable after averaging across 200 photostimuli (**Figure S1b,c**). The median threshold for PSP detection (PSP amplitude z-score ≤ –2.576, corresponding to a false-positive rate of 0.5%) was 0.35 mV (IQR 0.23 to 0.53 mV; n = 11) and was less than 1 mV in all cells assayed (**Figure 1g,h**). Optically measured PSPs were abolished upon pharmacological block of glutamatergic synaptic transmission (**Figure 1i**), confirming their synaptic origin. These results indicate that PSPs can be optically elicited from presynaptic terminals and demonstrate, for the first time, that they can be detected in postsynaptic neurons in a sensitive fashion.

To map presynaptic inputs to a population of motor cortical cells, we applied population-level optical connectivity measurements (**Figure 2a**) in the presence of tetrodotoxin (TTX) to prevent polysynaptic activation and 4-aminopyridine (4-AP) to enhance presynaptic excitability^33^, as in CRACM^16^. PSPs resulting from photoactivation of CheRiff+ axons from the contralateral motor cortex (“Contra”) were assayed across hundreds of cells in a single experiment (**Figure 2b**). In a single field-of-view, intermingled cells exhibited heterogeneous responses to axonal photoactivation (**Figure 2a,b and Figure S2**), with some showing large-amplitude PSPs and others responding minimally, or not at all (**Figure 2b and Figure S3**). Though GEVI photobleaching was negligible (**Figure S4**), in some cells, PSPs gradually decreased in amplitude during repetitive photostimulation in the presence of pharmacological blockers (**Figure S5**; see Supplementary Text). In these cells, we determined the number of trials across which averaging maximized the PSP signal-to-noise ratio (**Figure S6**). ‘Responding’ neurons had PSP amplitudes exceeding an SNR threshold (z-score) of –3.75, corresponding to a mean false-positive rate of 1.2% across all cells (**Figure S6a**). We estimated the minimum detectable PSP amplitude across all neurons at this SNR threshold, and found that it was < 1 mV (estimated) in 94% of cells (0.56 ± 0.29 mV, mean ± SD) indicating high PSP detection sensitivity (**Figure S6b**). Responding cells had a mean PSP amplitude of –0.93 ± 0.23% df/f (SEM; estimated 3.44 mV ± 0.85 mV), with a maximum PSP size of –6.72% df/f (estimated 25.89 mV), consistent with the amplitudes of optically elicited PSPs reported in the cortex in previous studies^30–32^.

**Figure 2.**
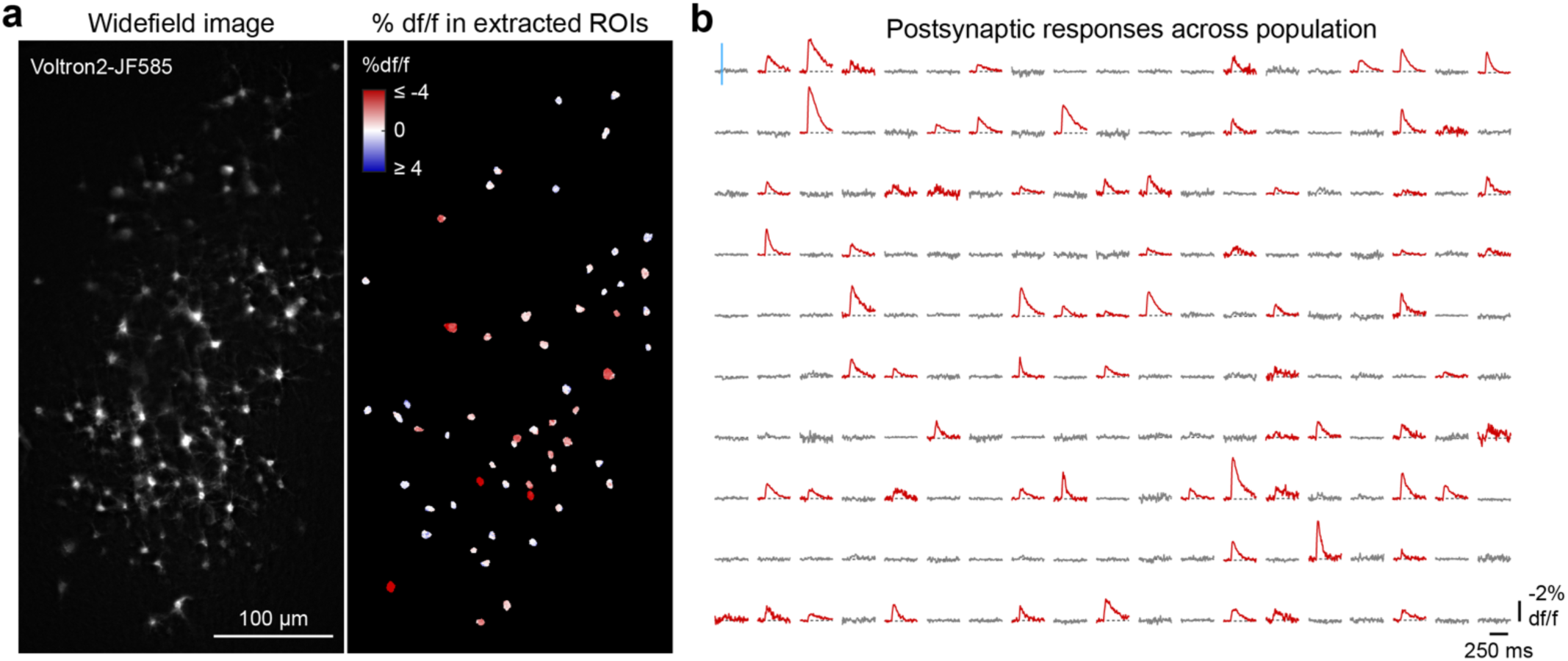
Population-level optical connectivity mapping. (**a**) Representative widefield field-of-view showing Voltron2-JF585 signal, spanning multiple cortical layers (left), with spatial heatmap of PSP amplitude (% df/f) measured for individual cell ROIs (right) in response to contralateral motor cortex axon activation. (***b***) PSP timeseries measured across imaged neurons from a single experiment, showing responding (red) and non-responding (grey) cells. Traces for a randomly selected 190 neurons are shown out of a total 218 imaged in one experiment.

These findings confirm that the presence and strength of long-range synaptic connections can be identified with high sensitivity using GEVI imaging, and that parallelizing detection dramatically increases the scalability of connectivity measurements compared to CRACM.

### Transcriptomic cell type identification in thick tissue sections

Spatial transcriptomic approaches that allow for the identification of many cell types often require thin (10-20µm) tissue sections or tissue expansion, and can necessitate specialized imaging systems^34–37^. We sought to develop an experimental and computational pipeline for transcriptomically identifying neurons interrogated with acute-slice GEVI imaging that can be performed routinely, at low cost, and in thick tissue sections without the use of gel-embedding or expansion. To achieve this, we optimized a protocol for multiplexed fluorescence in situ hybridization (mFISH) based on Hybridization Chain Reaction version three (HCRv3)^38^. This custom protocol permits high-SNR transcript detection throughout paraformaldehyde-fixed 300-µm-thick sections and across multiple rounds of probing/imaging using a standard confocal microscope. A series of permeabilization and clearing steps (**Figure S7a**) chemically quenched CheRiff-EGFP and JF585 fluorescence (**Figure S7b, Figure S8a**), and maintained high signal-to-background (**Figure S7c,d**) across multiple rounds of staining (**Figure S8b,c**). Using single-cell RNA sequencing (scRNA-seq) data from the Allen Brain Cell (ABC) atlas^11^, we selected a set of 17 genes whose expression differentiates motor cortical cell types (**Figure S9a**). This set incorporated marker genes commonly associated with both broad excitatory (*Rorb*, *Lratd2*, *Syt6*) and inhibitory (*Slc32a1*, *Pvalb*, *Sst, Vip*, *Lamp5*) cortical cell types, as well as additional differentially expressed genes that further distinguish excitatory types at a finer resolution (*Penk*, *Calb1*, *Pamr1*, *Dkkl1*, *Ccdc80*, *Myl4*, *Npnt*, *Slco2a1*; **Figure S9c**).

Next, we developed a computational pipeline for classifying cell types using multi-round mFISH image data. In each of 4 rounds of staining, transcripts associated with 4 marker genes were visualized (**Figure 3a**). A fifth pan-neuronal gene, *Snap25,* was included in all rounds of staining to ensure that RNA integrity was maintained across rounds. *Snap25* signal was also used for cross-round image registration (**Figure S8**) and for cell segmentation (**Figure S10**). Expression of each gene in mFISH data was evaluated in a binary fashion (i.e., expressed/not expressed) across cells to produce a 17-digit binary expression code for each segmented cell. To relate these codes to type-specific expression patterns in the ABC atlas cell type ontology^11^, we replaced each scRNA-seq cell with a “meta cell” representing its expression counts summed with the expression counts of its 9 nearest-neighbor cells of the same ABC “supertype” in PCA space to account for the relatively low RNA capture efficiency in droplet-based scRNA-seq data^39–41^. This process revealed bi-or trimodalities in marker gene expression that were not present at the single-cell level and which allowed for the selection of binary thresholds to demarcate cells “expressing” and “not expressing” each gene (**Figure S9b,c**).

**Figure 3.**
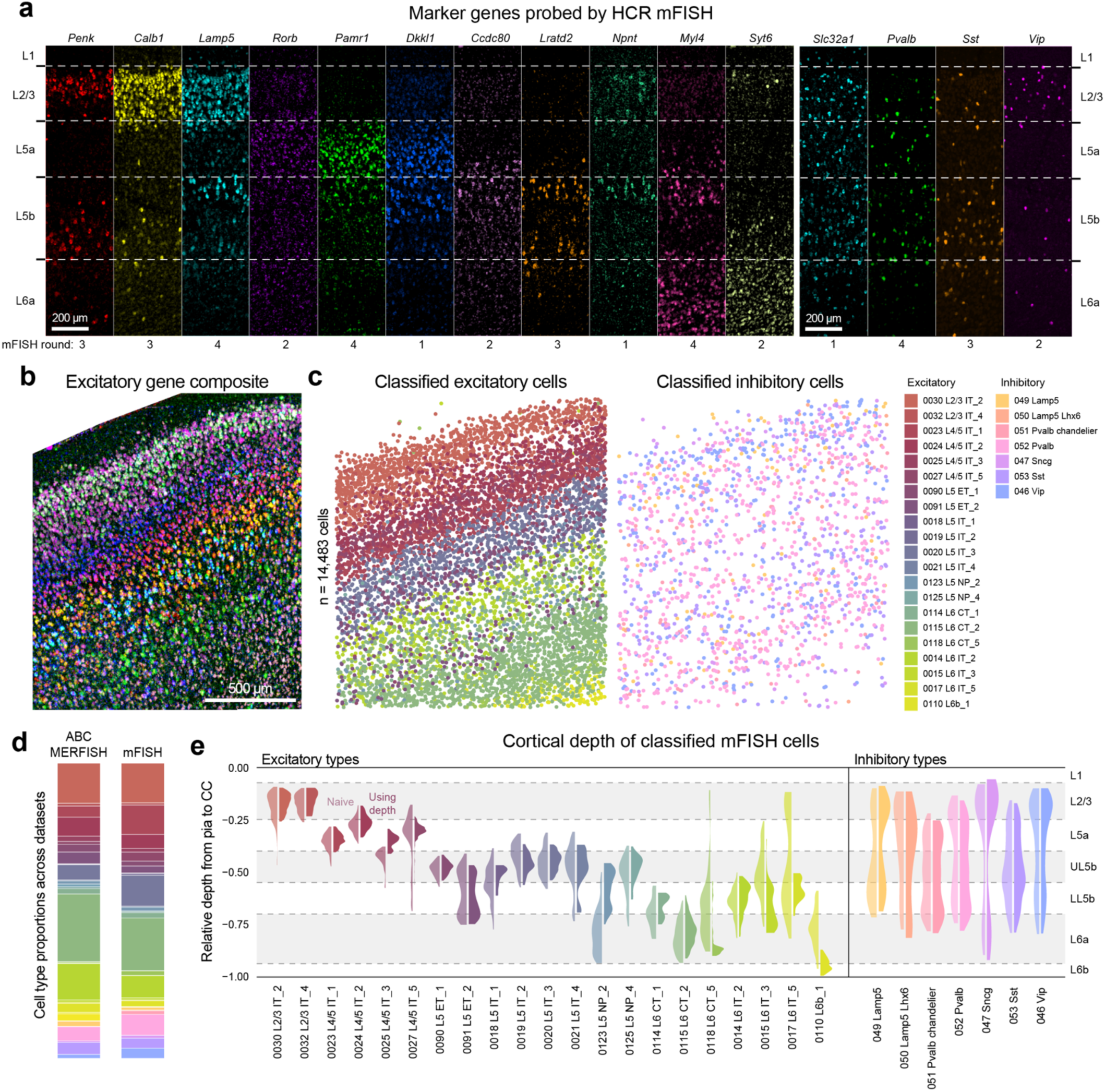
Cell type classification using thick-tissue mFISH data. (**a**) Confocal images of a subset of excitatory and inhibitory marker genes probed by HCR mFISH in one tissue section across 4 rounds of staining. (***b***) Z-projection and multichannel composite image of primary motor cortex showing mFISH signal for 12 excitatory marker genes across 3 confocal imaging planes. (***c***) Scatterplots showing motor cortex cells (dots) imaged across 50 µm of z-depth in the multi-round mFISH example shown in ***b*** and classified into excitatory (left) or inhibitory (right) cell types based on gene expression patterns (n = 14,483 cells). Cell types were determined using a classifier that included depth-dependent cell type priors and ABC scRNA-seq “meta cell” gene expression probabilities. (***d***) Stacked bar plots showing the cell-type composition of ABC MERFISH motor cortex data left) and the mFISH-based classification (right) for data shown in ***c***. (***e***) Violin plots of the laminar distribution of excitatory and inhibitory cell types classified from mFISH data shown in ***c*** and ***d***– without using (“naive”; left half of violin) or using (right half of violin) depth-dependent cell type priors. Distributions show cells between the 5^th^ and 95^th^ percentiles of depth.

We constructed a naïve Bayes classifier to assign cell type based on these binary gene expression patterns. The classifier included the proportion of every cell type that occupies each cortical depth (derived from ABC MERFISH data^11^) as a depth-dependent cell type prior (**Figure S11a-c**). We performed multi-round mFISH and applied this classifier to assign cell types at the ABC “supertype” level for excitatory cells and the broader “subclass” level for inhibitory cells (**Figure 3b-e**, **Figure S9d**; n = 14,483 cells). We found that cells assigned to each type largely followed expected laminar distributions whether depth information was included or not, with the exception of L6 intratelencephalic (IT) neurons whose classification accuracy was substantially improved through the inclusion of depth-dependent priors (**Figure 3e, Figure S11d**). The overall abundance and gene expression patterns of classified cells also largely matched those observed in ABC data (**Figure 3d, Figure S12**). Together, these results indicate that this pipeline can accurately and efficiently identify transcriptomic cell types in thick tissue.

### MOSAIX: A Multimodal Optical Strategy for Assaying Identity and Connectomics

We next integrated optical connectivity measurements with mFISH-based transcriptomic cell type identification to develop the full MOSAIX pipeline (**Figure 4a, Figure S13**) and determine how long-range inputs are integrated in the motor cortex in a cell-type specific manner. We first determined whether Voltron2 could be expressed broadly across all cell types via AAV injection. As expected, cell-type tropisms were serotype-dependent (**Figure 4b-d**). We found that delivering Cre via AAV5 yielded broad Voltron2 expression across cell types (**Figure 4d**) and selected this serotype for subsequent experiments. We then expressed Voltron2 in motor cortex cells and CheRiff in either MThal (**Figure S14a**) or Contra (**Figure S14b**) – two of the most prominent sources of afferent input to motor cortical neurons. The distribution of axonal CheRiff across cortical layers was similar following MThal or Contra CheRiff expression (**Figure S14c,d**), with the notable exception of layer 1, which contained a high density of MThal, but not Contra, axonal labeling. We then applied MOSAIX to identify motor cortex neurons receiving synaptic connections from MThal (n = 2 animals; **Figure 4e**) and Contra axons (n = 2 animals; **Figure 4f**).

**Figure 4.**
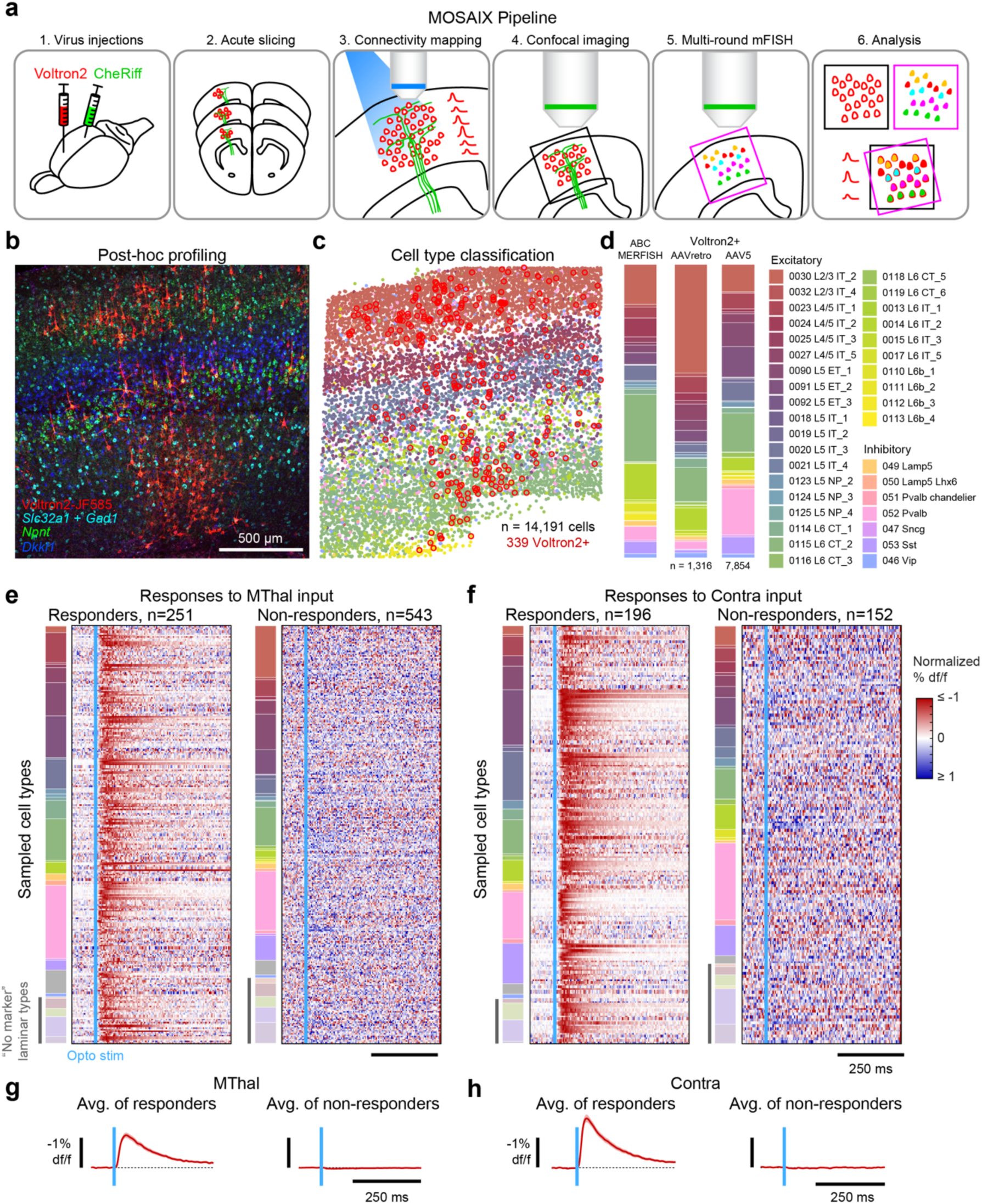
Combined MOSAIX pipeline provides cell-type-specific connectivity data. (**a**) Schematic outline of complete MOSAIX pipeline. (**b**) Confocal image of HCR mFISH performed in Voltron2+ motor cortex. (***c***) Scatterplot showing motor cortex cells (dots) imaged across 50 µm of z-depth in multi-round mFISH and classified into excitatory or inhibitory cell types (n = 14,191 cells). Voltron2+ cells (n = 339 cells) are highlighted with red circles. (***d***) Stacked bar plots showing the cell-type composition of ABC MERFISH data (left), and the Voltron2-expressing population when the Cre transgene is delivered using either the AAVretro (center, n = 2 animals) or AAV5 (right, n = 4 animals) serotype. (***e, f***) Heatmaps showing PSP timeseries normalized to the maximum value in the response window across individual responding (left) and non-responding (right) cells (rows) imaged in MOSAIX experiments mapping MThal input (***e***; n = 2 animals, 794 cells) or contra motor cortex input (***f***; n = 2 animals, 348 cells) onto motor cortex. Rows ordered first by cell type (highlighted in stacked bar to the left) and then by PSP amplitude. Cells with no detectable mFISH markers (i.e., “No marker” laminar types) highlighted with grey bar. (***g, h***) Average PSP profiles across all responding (left) and non-responding (right) cells in the MThal circuit (***g***) and the Contra circuit (***h***). Shaded region shows SEM.

Following optical connectivity measurements, brain slices were fixed and high-resolution confocal images were acquired to verify that Voltron2+ neurons were located within the terminal zone of CheRiff+ axons and to facilitate registration of GEVI imaging with post hoc mFISH labeling (**Figure S15, Figure S16a**). Gene expression was then evaluated using multi-round mFISH. Cells were segmented based on *Snap25* signal, as before (**Figure S10b,c**), but only Voltron2+ cells were further analyzed (**Figure S16b**). A subset of Voltron2+ cells (18%) lacked detectable mRNA (possibly due to cellular stress from viral transduction or mechanical damage) and were therefore excluded from subsequent analyses. Using MOSAIX, the connectivity and transcriptomic identity of 794 (MThal) and 344 (Contra) neurons was identified (**Figure 4e,f**). Importantly, the sampled populations showed similar cell type composition across both circuits, with all excitatory supertypes and inhibitory subclasses represented in both datasets.

Postsynaptic potentials were detected in 251/794 cortical cells (31.5%) upon MThal axon photoactivation and 196/344 cortical cells (57.0%) upon Contra axon photoactivation (**Figure 4e,f**), with responder proportions varying across cell types (**Figure S17**). Across individual experiments assaying the same circuit, mean cell type PSPs varied in amplitude (**Figure S18**), likely due to differences in opsin expression (MThal #1: –0.25 ± 0.05%, mean ± SEM; MThal #2: –0.35 ± 0.07%; Contra #1: –0.46 ± 0.13%; Contra #2: –1.35 ± 0.28%). In order to account for this variation, PSPs were normalized to the mean amplitude of the cell type with the largest response, so that the mean response amplitude of all cell types was between 0 and 1.

Responses to MThal and Contra input were strongest in cells from layer 5a to layer 6a (L5a and L6a; **Figure 5a,b, Figure S19, Figure S20**), closely resembling known motor thalamic and corticocortical innervation patterns^42,43^. To further dissect the cell-type specificity underlying these connectivity patterns, we compared PSP amplitudes in ABC “supertypes” comprising the three most prominent excitatory cell classes in layers 5 and 6: intratelencephalic (IT), extratelencephalic (ET), and corticothalamic (CT) cells (**Figure 5c-e**). Interestingly, IT supertypes in L5 showed a high degree of heterogeneity in their responses to both MThal and Contra input. The *Rorb*-“0020 L5 IT_3_” supertype exhibited PSP amplitudes 2.17x larger than *Rorb*+/*Dkkl1*-“0027 L4/5 IT_5_” cells and 1.79x larger than *Rorb*+/*Dkkl1*+ “0023 L4/5 IT_1_” cells following photoactivation of MThal axons (L4/5 IT_5_: 0.46 ± 0.25, n = 27; L4/5 IT_1_: 0.56 ± 0.20, n = 40; L5 IT_3_: 1.00 ± 0.45, n = 37; p < 0.0001, one-way ANOVA). Similarly, activation of Contra axons produced PSPs in L5 IT_3_ cells that were 3.14x larger than in L4/5 IT_5_ cells and 2.44x larger than in L4/5 IT_1_ cells (L4/5 IT_5_: 0.21 ± 0.16, n = 9; L4/5 IT_1_: 0.27 ± 0.17, n = 14; L5 IT_3_: 0.66 ± 0.25, n = 30; p < 0.0001, one-way ANOVA).

**Figure 5.**
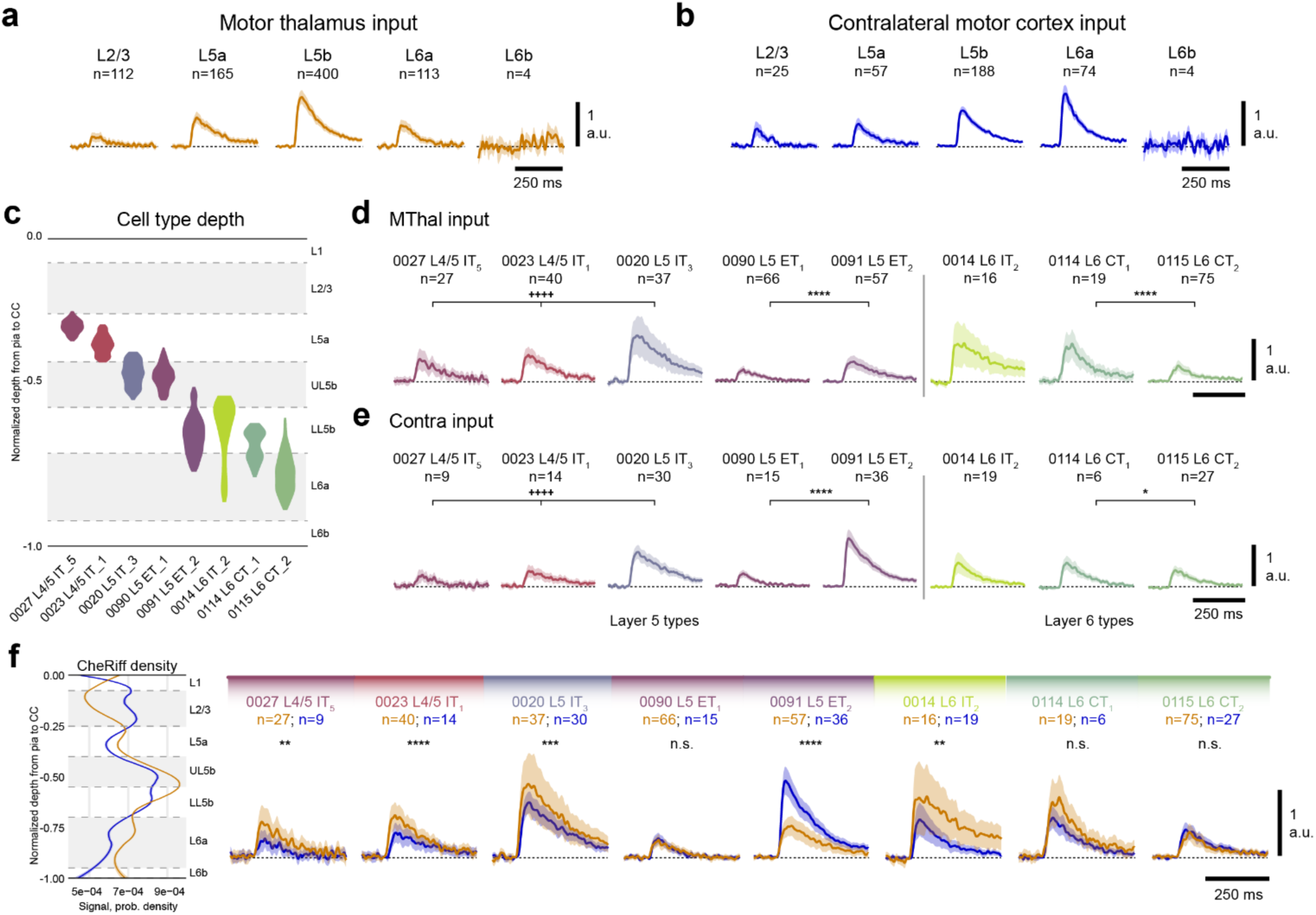
MOSAIX provides cell-type-specific insights into long-range circuit connectivity. (**a, b**) Scaled and averaged motor cortex PSPs (averaged across both“responding” and “non-responding” cells) resulting from activation of MThal inputs (***a***), or Contra inputs (***b***), grouped by laminar position. Line shows mean PSP trace, shaded area shows SEM. (***c***) Violin plots of the laminar position of the ABC-derived cell “supertypes” shown in (***d-f***). (***d, e***) Scaled and averaged PSPs for each supertype resulting from MThal input (***d***) or Contra input (***e***). Shaded region shows SEM; **^++++^** = p < 0.0001 by one-way ANOVA; **** = un-adjusted p < 0.0001, n.s. = not significant by two-tailed t-test. (***f***) Probability density distributions (left) of CheRiff-EGFP signal across all MOSAIX imaged sections in the MThal-to-motor cortex circuit (orange), and the Contra-to-motor cortex circuit (blue; line shows mean, shaded area shows 95% CI), alongside scaled and averaged PSPs of cell supertypes shown in ***d*** and ***e*** (right), colored based on activated pathway (MThal in orange, Contra in blue; line shows mean PSP trace, shaded area shows SEM; *** = p < 0.001, **** = p < 0.0001; n.s. = not significant). All vertical scale bars show the maximum average value (1) used for scaling responses within each experiment.

Importantly, normalizing each cell’s PSP amplitude to CheRiff intensity within its cortical column maintained relative differences in response strength across cell types (**Figure S21**), indicating that spatial variation in innervation density did not skew results. Transcriptomically distinct IT neurons possess diverging projection patterns^44^, suggesting that thalamic and callosal signals to motor cortex may be preferentially relayed to specific downstream cortical and striatal areas.

Across ET supertypes, we observed that both MThal and Contra input targeted *Npnt*-/*Slco2a1*+ “0091 L5 ET_2_” neurons much more strongly (by 2.00x and 4.35x, respectively) than *Npnt*+/*Slco2a1*-“0090 L5 ET_1_” neurons (MThal – ET_1_: 0.21 ± 0.10, n = 66; ET_2_: 0.42 ± 0.15, n = 57, p < 0.0001, two-tailed t-test; Contra – ET_1_: 0.23 ± 0.09, n = 15; ET_2_: 1.00 ± 0.26, n = 36; p < 0.0001; **Figure 5d,e**). This high degree of specificity is surprising given the overall transcriptomic similarity of ET_1_ and ET_2_ neurons. Medulla-and spinal-cord-projecting cortical neurons belong to the ET_2_ supertype, and the strong afferent innervation of these neurons could account for their strong modulation during movement. In contrast, ET_1_ neurons, which target the thalamus and avoid motor circuits in the medulla and spinal cord, are more weakly modulated during movement onset and appear to receive much weaker afferent inputs^45^.

Lastly, across CT supertypes, we found that MThal weakly targeted *Lamp5*-“0115 L6 CT_2_” cells, which represent the large majority of CT neurons (**Figure 5d,e**). This is consistent with previous findings that thalamic input largely avoids thalamus-projecting L6 neurons^31,42,46^. However, MOSAIX also revealed that rarer *Lamp5*+ “0114 L6 CT_1_” cells, many of which are located in lower L5b, are more potently innervated by MThal and Contra inputs, with MThal innervation eliciting PSPs 2.52x larger (CT_1_: 0.68 ± 0.38, n = 19; CT_2_: 0.27 ± 0.14, n = 75; p < 0.0001, two-tailed t-test). This could indicate the existence of a monosynaptic thalamo-cortico-thalamic loop mediated by a relatively rare CT type. Contra input strength to this CT_1_ type was 1.41x stronger than to the CT_2_ type (CT_1_: 0.51 ± 0.18, n = 6; CT_2_: 0.36 ± 0.13, n = 27; p = 0.0163), indicating that the CT_1_ type receives greater afferent input across both circuits.

We next examined the relative innervation of cortical cell types across the MThal and Contra pathways (**Figure 5f**). Interestingly, the ratio of input from these sources was similar across all IT cell types in layers 4/5a/5b/6a, with MThal input appearing modestly larger when responses were normalized per experiment (p < 0.01, two-tailed t-test). ET_2_ cells deviated markedly from this pattern, showing greater response amplitudes following Contra activation compared to MThal input (p < 0.0001, two-tailed t-test). These differences were observed despite an overall similarity in the laminar profile of axonal labeling across these pathways, suggesting that afferent connectivity patterns are not a simple function of laminar axon density.

Together, our findings suggest that afferent inputs to the motor cortex establish synaptic connections in a manner that is highly specific, even for cell types belonging to the same broad class (i.e. IT, ET, CT, etc.). Determining the architecture of afferent cortical connectivity at the level of precisely defined cell types has not been possible with existing methods. The integration of scalable methods for measuring synaptic connectivity and assigning transcriptomic identity in MOSAIX now allows cell-type specific connectivity motifs to be characterized efficiently in any pathway in the brain.

## Discussion

We describe MOSAIX, an integrated platform for measuring the presence and strength of long-range synaptic connections onto many transcriptomically identified neurons. To measure synaptic connectivity, MOSAIX integrates optogenetic control of afferent axons and optical detection of PSPs using voltage imaging. This approach for optical connectivity is highly parallelizable, and we demonstrate here, for the first time, that it can be used to detect postsynaptic potentials across large populations with sub-millivolt sensitivity. Importantly, we demonstrate that high-throughput connectivity measurements can be readily combined with spatial transcriptomics, a scalable approach for cell type identification. Leveraging MOSAIX, we assayed the strength of thalamic and contralateral synaptic input onto more than 1,000 transcriptomically identified neurons in the mouse primary motor cortex. Cell-type-specific measurements of afferent connectivity have never before been demonstrated at this scale. With MOSAIX, large-scale measurements could be conducted efficiently, requiring only a small number of experiments.

Examining motor thalamic (MThal) and contralateral (Contra) inputs to the motor cortex allowed us to validate MOSAIX through comparisons with prior studies examining afferent inputs from these sources, either by lamina, or onto broadly defined cell types^42,43,46^. MOSAIX recapitulated known patterns of thalamocortical and cortico-cortical connections to the motor cortex, such as the strong innervation of L5 cells by both pathways (**Figure 5a**), establishing its fidelity.

The scalability of MOSAIX, and the ability to classify cell types precisely, revealed several novel cell-type-specific afferent connectivity motifs. For example, we found that the *Dkkl1*+/*Ccdc80*+ IT supertype in upper L5b is driven powerfully compared to other L5 IT cell types, and the *Slco2a1*+ ET_2_ supertype is much more strongly innervated by long-range sources compared to the neighboring, and transcriptomically similar, *Npnt*+ ET_1_ supertype (**Figure 5d,e**). The latter observation suggests that the fast modulation of ET_2_ dynamics during movements^45^ may be driven by inputs originating outside of the motor cortex – an intriguing new hypothesis that can be examined more closely in future work. Further, MOSAIX revealed strong input to a superficial *Lamp5*+ CT type, particularly through the MThal pathway, compared to the deeper and more prevalent *Lamp5*-CT type. Previous work had suggested a paucity of thalamic inputs to CT neurons, indicating the absence of a strong monosynaptic thalamo-cortico-thalamic connectivity mediated by CT neurons. Our results suggest that such a circuit motif may indeed exist, mediated by a comparatively rare CT type that is displaced into L5b.

Together, our findings establish that cell types belonging to the same broad classes and located at similar cortical depths are often innervated in a highly disparate manner. This observation underscores the critical importance of high-resolution cell type identification, which has not been possible in the large majority of prior studies of synaptic connectivity motifs using CRACM or other methods.

MOSAIX builds upon previous work that leverages optical strategies to map synaptic connectivity. In “Optomapping”, opsin-expressing presynaptic neurons are serially activated optically while resulting PSPs are detected in a single neuron with whole-cell electrophysiology^47^. “SynOptopatch” also leverages GEVI imaging and optogenetics to measure synaptic responses across populations of neurons^17^, although whether it achieves sufficient sensitivity to detect PSPs reliably is not clear. “Voltage-Seq” also employs optogenetic activation of afferent axons and postsynaptic voltage imaging using a similar approach, but has principally been used to examine network dynamics engaged by long-range inputs – an application that does not require the sensitive detection of monosynaptic PSPs^48^. Cell type identification in these methods is achieved using transgenic animals (SynOptopatch) that can restrict GEVI expression to a single cell type, or pipette-based extraction of individual imaged cells for RNA-sequencing (Voltage-Seq). Both of these approaches suffer from low-throughput in cell identification, and therefore cannot be utilized to assign transcriptomic type precisely at scale.

There are clear opportunities for further improvement of the scalability and sensitivity with which connections are identified – and the precision with which cell types can be classified – in MOSAIX. Using advanced imaging methods that provide optical sectioning^49,50^ would permit substantially denser GEVI expression and yield commensurately higher throughput. Sensitivity can also be improved by applying optical sectioning to reduce background fluorescence, or through further GEVI optimization^51^. Such optimizations are likely to permit unitary synaptic potentials (typically 0.1 to 2.5 mV in amplitude in the cortex^52–54^) to be reliably detected. To discern cell types with a higher degree of precision, high-plex spatial transcriptomics methods^55^ can be used in place of mFISH to provide greater transcriptomic resolution than we present here. Such methods are advancing rapidly.

A critical component of this work is the demonstration that high PSP detection sensitivity can be achieved across populations of neurons using voltage imaging. This provides a number of exciting opportunities, for example, modest improvements to PSP detection sensitivity, in conjunction with single-neuron photostimulation (as in Optomapping^47^), would enable individual synaptic connections between pairs of neurons to be measured optically with high fidelity. Such a paradigm has the potential for even greater scalability, as the number of pairwise synaptic connections that can be assayed simultaneously scales with the square of the number of GEVI-labeled neurons imaged within a field-of-view. This could yield an ultra-high-throughput alternative to paired-patch electrophysiology for measuring synaptic connections, where the connectivity between many thousands of cell pairs could be investigated in minutes.

The development of MOSAIX, a highly parallelizable, all-optical approach for quantifying synaptic connectivity, provides a novel, scalable approach for determining the connectivity motifs that define the architecture of neural circuits containing many cell types. High-throughput methods for measuring connectivity will pave the way for examining how circuits vary across individuals and how they are remodeled as a result of learning or disease.

## Resource availability

All data and code will be made fully available on Zenodo.org

## Acknowledgements

We thank A. Abdelfattah and J. Mertz for advice and discussions, the Boston University Micro/Nano Imaging Facility for providing imaging resources, and the University of Pennsylvania Vector Core for production of the CheRiff-encoding AAV. We additionally thank A.M. Zambon for assistance in the lab, and J. Chen, T. Wang, and N. Li for comments on the manuscript. This work was supported by research grants from the National Institutes of Health: RF1MH126882 (MNE), F32MH129149 (MVM), and K99NS139312 (MVM). The funder had no role in study design, data collection and analysis, decision to publish, or preparation of the manuscript.

## Author Contributions

MVM and MNE conceived of the project. MNE supervised research. MVM performed experiments, with assistance from JPV, and with resources provided by TW. MVM and WJC analyzed data. MVM and MNE wrote the paper.

## Declaration of interests

The authors declare no competing interests.

## Supplemental Information

Document S1. Supplemental text and Figures S1-S21

## STAR Methods

### EXPERIMENTAL MODEL AND SUBJECT DETAILS

#### Animals

All animal procedures and experiments were performed with approval from the Boston University Institutional Animal Care and Use Committee (IACUC) and in compliance with policies and guidelines from the National Institutes of Health (NIH). Both male and female C57BL/6J mice (JAX #000664; RRID: IMSR_JAX:000664) were used for all experiments described. Mice were housed on a 12-h light/dark cycle at 21 ± 3°C and 30–70% humidity with ad libitum access to food and water.

### METHOD DETAILS

#### Virus injections

Virus was injected into animals between 5-7 weeks of age. Anesthesia was induced using 4% isoflurane until animal was unresponsive to toe pinch. Animals were then maintained under anesthesia using a constant flow of 0.8% isoflurane through a nose cone in the stereotaxic apparatus. Saline ointment was applied to both eyes, and pre-operative Ketoprofen (4 mg/kg, 1 mg/mL solution in sterile saline) and Buprenorphine HCl (0.072 mg/kg, 0.03 mg/mL solution in sterile saline) analgesics were delivered via subcutaneous injection (S.C.). Bupivacaine HCl (100 µL of a 5 mg/mL solution in sterile saline) was delivered subcutaneously to numb the scalp and periosteum. Nair hair removal cream was applied to the scalp to remove fur. Betadine was then topically applied and the scalp and periosteum were resected to expose the skull. Exposed scalp margins were secured to the skull with cyanoacrylate glue. The animal’s head was oriented to bring the bregma and lambda sutures to the same z-depth, and to ensure the bilateral parietal skull plates were leveled. Craniotomies were then made over target regions of the brain, leaving only a thin layer of bone that could be punctured by a beveled pipette. To express CheRiff-EGFP in presynaptic neurons, 50 nL of a sterile PBS solution containing a custom AAV9-hSyn-CheRiff-EGFP virus (titer of 1.0E12 g.c.) was delivered via glass pipette to motor thalamus (MThal) at –1.6 mm AP, +0.8 mm ML, –4.25 mm (VM of MThal) and –1.0 mm AP, +1.3 mm ML, –3.6 mm DV (spanning AM, VA, and VL of MThal; coordinates from bregma), or to contralateral motor cortex (Contra) at 6 different AP/ML sites located at +0.7/1.3/1.9 mm AP, –1.9/1.5 mm ML, –1.0 mm and –0.5 mm DV. In ipsilateral motor cortex, 50 nL of sterile PBS solution containing AAV1-hSyn-FLEX-Voltron2 (titer of 1.0E12 g.c.) mixed with AAVretro-hSyn-Cre or AAV5-hSyn-Cre (titer of 0.5-1.0E11 g.c.) was delivered to two depths at 3 different AP sites: +0.7/1.3/1.9 mm AP, +1.7 mm ML, –1.0 mm and –0.5 mm DV. Virus solutions were delivered at a rate of about 1 nL/sec using a Narishige microvolume ejector. The skull of the animal was then covered with dental cement to cover surgical areas. Animals were given postoperative Ketoprofen once per day for the following two days. Viral construct expression was allowed to carry on for 3-4 weeks.

#### Acute slicing

Acute slices were made using an N-methyl-D-glucamine (NMDG)-based protective recovery method^56^. Briefly, animals were perfused with 15 mL chilled and carbogen-bubbled (95% O2/5% CO2) NMDG-HEPES aCSF (in mM: 92 NMDG, 2.5 KCl, 1.25 NaH2PO4•2H2O, 30 NaHCO3, 20 HEPES, 25 D-Glucose, 2 Thiourea, 5 (+)-Na-L-ascorbate, 3 Na-pyruvate, 0.5 CaCl2•2H2O, 10 MgSO4•7H2O, pH adjusted to 7.3-7.4 with HCl) at 10 mL/min. The brain was then rapidly dissected in chilled and bubbled NMDG-HEPES aCSF and mounted for vibratome sectioning. 300 µm slices were made in a para-coronal plane (with anterior tissues oriented slightly ventrally) through motor cortex with chilled and bubbled NMDG-HEPES in the bath. Additional acute slices were collected through MThal injection sites if present, immediately fixed in fresh 4% paraformaldehyde in PBS for 30 min, and stored at 4°C in PBS. Motor cortex slices were immediately moved to a chamber with 34°C bubbled NMDG-HEPES, where NaCl was reintroduced gradually via periodic spike-ins of 2M NaCl in NMDG-HEPES at 5 min. intervals. Slices were then incubated for 1 hour in a small chamber containing 25 nmol of JF585 dye in room temperature, bubbled HEPES aCSF (in mM: 92 NaCl, 2.5 KCl, 1.25 NaH2PO4•2H2O, 30 NaHCO3, 20 HEPES, 25 D-Glucose, 2 Thiourea, 5 (+)-Na-L-ascorbate, 3 Na-pyruvate, 2 CaCl2•2H2O, 2 MgSO4•7H2O, pH adjusted to 7.3-7.4 with NaOH). Finally, slices were transferred to a chamber containing room temperature, bubbled HEPES aCSF, where they were held until imaging. Imaging experiments were performed in either the above HEPES aCSF or in recording aCSF (in mM: 124 NaCl, 2.5 KCl, 1.2 NaH2PO4•2H2O, 24 NaHCO3, 5 HEPES, 12.5 D-Glucose, 2 CaCl2•2H2O, 2 MgSO4•7H2O, pH adjusted to 7.3-7.4 with NaOH). MOSAIX experiments were performed with 200nM-1µM TTX and 100 µM 4-AP (Tocris #1069 and #0940) in the extracellular recording aCSF. For experiments including glutamatergic blockers, 10 µM NBQX and 10 µM D-AP5 (Tocris #1044 and #0106) were applied to the extracellular recording aCSF.

#### Whole-cell patch clamp

For whole-cell recording experiments, fire-polished, filamented glass pipettes (1.5 mm OD, 0.86 mm ID, #BF150-86-10 Sutter Instruments) were pulled to 3-7 MΩ resistance using a P-1000 horizontal micropipette puller (Sutter Instruments). Pipettes were loaded with an internal solution containing (in mM) 130 K-gluconate, 4 KCl, 10 HEPES, 0.3 EGTA, 10 Phosphocreatine-Na2, 4 MgATP, and 0.3 Na2GTP. Cells were visualized using oblique illumination with an M660L4 LED (ThorLabs) through a condenser focused through the glass of the bath onto the sample to produce a DIC-like image. An Axon Instruments MultiClamp 700b amplifier and patch-clamp headstage (Molecular Devices #1-CV-7B) mounted on a motorized four-axis Siskiyou MX7600 manipulator were used to acquire all electrophysiology data.

#### Voltage imaging and optogenetic photoactivation

During voltage imaging, slices were continually perfused with room temperature, bubbled HEPES aCSF via peristaltic pump. MOSAIX experiments were performed with 200 nM-1 µM TTX and 100 µM 4-AP added to the perfused aCSF. Widefield images were acquired using a Nikon 16X/0.8NA water-immersion objective (#CFI75 LWD 16X W) or an Olympus 20X/1.0NA water-immersion objective (#N20X-PFH). To illuminate the sample, a M565L3 LED (Thorlabs) was used with an ET 577/25x excitation filter, T590lpxr dichroic beam splitter, and an ET 590lp emission filter (Chroma). A 300mm 200 EFL lens (Edmund optics) was used to focus the signal onto the sensor of an ORCA-Flash4.0 v3 CMOS (Hamamatsu, model #C13440-20CU) or a Kinetix sCMOS (Teledyne Vision Solutions, model #01-KINETIX-M-C). For optogenetic photoactivation, an M470L4 LED was used. During MOSAIX experiments, LEDs and camera acquisition was triggered using digital waveforms generated in Wavesurfer (https://wavesurfer.janelia.org/) via a digital-to-analog converter (National Instruments #BNC 2090), and LED Drivers (Thorlabs #LEDD1B). The 565nm imaging LED was on continuously for all acquisitions (approximately 25 mW/mm^2^). For whole-cell patch experiments, the 470nm opto stim LED was pointed obliquely at the sample, achieving approximately 0.1 mW/mm^2^ intensity which could elicit small amplitude responses for PSP detection sensitivity measurements. Stim pulse duration was adjusted to modulate response amplitude, with pulse widths ranging from 0.5 to 5.0 ms. For MOSAIX experiments, the opto stim LED was used at approximately 8 mW/mm^2^ (1.0 ms pulse width) and placed below the sample, with light focused onto the sample through a condenser. Opto stim was performed at 2 or 4 Hz for patch clamp experiments and 1 or 2 Hz for MOSAIX experiments, while continuously acquiring Voltron2-JF585 fluorescence signal at 400 Hz. For patch clamp experiments, either a 128 x 128 pixel or 1024 x 256 pixel field-of-view (FOV) with a 2x binning factor was used to acquire data. For MOSAIX experiments, variable FOV sizes were used, aiming to image as many cells per section as possible while maintaining a 400 Hz imaging rate. After completion of MOSAIX, imaged sections were incubated in 4% paraformaldehyde in RNase-free PBS for 30-45 minutes at room temperature (RT) with gentle agitation, then stored in PBS at 4°C.

#### Pre-HCR mFISH histology

MOSAIX sections were microdissected to remove all but the dorsal cortical region containing Voltron2-positive neurons in motor cortex, then incubated for 1.5-2 hrs at RT in a dye mixture containing 1:1000 NeuroTrace 435/455 and 1:100 SYTO 45 dye (Thermo Fisher #N21479 and #S11350 respectively) in PBS to label nucleic acids in all neurons in the sample. motor cortex sections were briefly washed and mounted in 85% 2,2’-thiodiethanol (TDE) in ultra-pure water, ensuring that the MOSAIX-imaged slice face was face up on the slide. NeuroTrace 435/SYTO 45, CheRiff-EGFP, and Voltron2-JF585 fluorescence was imaged on a Nikon CSU-W1 SoRa spinning disk confocal microscope. 40X image stacks with 2 µm spacing through 50-80 µm of axial z-depth were collected, with enough XY tiled sampling to visualize all Voltron2-positive neurons in each section. Sections were removed from the slide and washed for 15 min in PBS at RT before proceeding to our custom HCR mFISH protocol.

#### Multi-round mFISH protocol for thick tissue

Multiplexed FISH was performed using an adapted version of the Molecular Instruments HCRv3 “generic sample in solution” protocol^38^, incorporating multi-round probing and stripping^37,57^. All chemical reagents were not used for more than 2 months after opening, and all solutions were prepared using nuclease free stocks, salts, water. RNaseZap (Thermo Fisher #AM9780) or RNase AWAY (Thermo Fisher #7002) was used to clean all instruments used during the HCR procedure. Unless otherwise stated, all washes and incubations were performed with end-over-end rotation in 600 µL microcentrifuge tubes, using 200 µL of solution per slice and per tube. Day 0: After pre-HCR imaging, MOSAIX samples were incubated in 90% DMSO in PBS at RT for 2 hours. After 1 brief rinse in PBS, slices were each moved to a well of a 12-well plate containing freshly prepared 1% NaBH4 in PBS and incubated at RT for 15 minutes on an orbital shaker. Slices were carefully transferred back to a new microcentrifuge tube containing PBS and washed 3 x 5 min at RT to remove all NaBH4 solution. Slices were stored in fresh PBS at 4°C until the following day. Day 1: PBS from the sample tubes was exchanged for 8% SDS in PBS (prepared fresh from a 10% SDS stock in water) and slices were incubated for 4 hours at RT. Excess SDS was removed by washing the slices 3 x 5 min and then 3 x 1 hr in PBS at RT. From this point, the Molecular Instruments HCRv3 protocol for staining a “generic sample in solution” was followed, using the reaction volumes listed above. Briefly, slices were pre-incubated in Hybridization buffer (Molecular Instruments) for 30 min at 37°C. An initiator probe solution containing a final concentration of 4 nM of each initiator probe set (ie., 4 nM per gene of interest) was made in Hybridization buffer. The pre-incubation solution was then exchanged for this initiator probe solution and samples were incubated overnight (ON) at 37°C. Day 2: Wash buffer (Molecular Instruments) was pre-warmed to 37°C by placing into the incubator for approximately 15 minutes before using. Slices were washed in warmed Wash buffer 2 x 15 min and then 2 x 30 min at 37°C. Slices were then washed in 5X SSC with 0.1% Tween20 (5X SSCT) 3 x 5 min at RT, and then stored at 4°C until the end of the day. To end the day, samples were pre-incubated in Amplification buffer (Molecular Instruments) for 30 min at RT. During this time, each set of amplifiers needed (per 200 µL reaction volume, 2 µL of h1 and 2 µL of h2 were used per amplifier pair) was melted separately by heating to 95°C for 90 sec and then cooled to room-temperature by 5° per minute in a thermocycler. Amplifiers were spun down and then pooled in fresh Amplification buffer. The pre-incubation solution was then exchanged for the amplifier solution and slices were incubated ON at RT, protected from light. Day 3: Samples were washed 3 x 5 min in 5X SSC (no Tween20) and then incubated in 85% TDE in water to clear before mounting. Slices were carefully moved from the tube to a glass slide and coverslipped in sufficient 85% TDE to cover the samples. Slices were imaged on an Olympus FV3000 laser scanning or Nikon CSU-W1 SoRa spinning disk confocal microscope. 20X or 40X image stacks with 2 µm spacing through 50-80 µm of axial z-depth were collected, with enough XY tiled sampling to visualize the full area of motor cortex that contained Voltron2-positive neurons in each section. Following imaging, the coverglass was removed from the slide, slices were rehydrated with 2X SSC and carefully lifted from the glass with a brush. Samples were placed back into fresh microcentrifuge tubes containing 2X SSC buffer and washed 3 x 5 min in 2X SSC to remove excess TDE. Slices were then pre-incubated in 1X DNaseI buffer (New England Biolabs #B0303) at 37°C for 15 min before being incubated in 1:50 DNaseI (enzyme stock of 2000 U/mL, New England Biolabs #M0303; working concentration of 8 U per 200 µL reaction volume) in 1X DNaseI buffer for 1 hr at 37°C. This solution was then exchanged for a fresh solution of 1:50 DNaseI in 1X DNaseI buffer, and slices were incubated ON at 37°C. Day 4: Samples were incubated in 65% formamide in 2X SSC for 30 min at 37°C. After 2 brief rinses in 2X SSC, each slice was then carefully transferred to a well of a clean well-plate containing freshly prepared 1% NaBH4 in PBS, and incubated for 15 min at RT on an orbital shaker. Slices were then moved back to a clean microcentrifuge tube containing PBS, and washed 3 x 5 min in PBS at RT. Slices were then stored in PBS at 4°C until the end of the day, when the second round of initiator probe hybridization was performed, as described for the end of Day 1. From here, steps repeat as described above until 4-5 rounds of HCR have been performed.

#### Voltage imaging data processing and analysis

All voltage imaging analyses were performed in MATLAB (Mathworks). Motion correction was performed on widefield timeseries images of Voltron2-JF585 signal using a modified, GPU-accelerated implementation of the NoRMCorre algorithm^58^. Rigid registration was applied frame-by-frame using a subpixel-accurate template-matching approach. A representative template was constructed from a batch of 200 frames starting at frame 800 (or the midpoint of the movie, if shorter), and all frames were aligned to this reference using a maximum shift of 15 pixels and an up-sampling factor of 50. To correct for large-spatial-scale intensity variations, background was removed using a custom implementation of the rolling ball algorithm, based on the method used in FIJI’s Subtract Background plugin^59^. For each frame, a local background estimate was computed by performing a morphological opening on a lightly blurred image with a spherical structuring element of radius larger expected foreground structures. This background was then subtracted from the original frame. Processing was done frame-by-frame to preserve neural activity that varied on faster timescales than the background, ensuring high-frequency signal components were retained. Timeseries data for all imaged cells were extracted from each image frame as the sum of the total signal within the area of each cell’s ROI (drawn manually for whole-cell patch experiments, or derived from mFISH segmentation masks for MOSAIX experiments – see below). Percent changes in fluorescence over the median baseline signal (% df/f) were then calculated for each cell. As before, timeseries were segmented based on the frequency of photostimulation to isolate each photoactivation /PSP instance, including a post-response baseline period (where cell was at resting). These individual PSP instances were averaged according to the “optimal” number of trials determined for each cell (described below). For replicate MOSAIX experiments, the cell-type-specific PSPs were scaled before averaging across individual experiments to account for potential differences in CheRiff excitation. Responses were scaled to the mean PSP amplitude of the cell type with the largest response in each experiment, ensuring that all mean PSP amplitudes fell between 0 and 1.

#### Voltage imaging sensitivity evaluation

Both whole-cell recording and voltage imaging were segmented in time based on the frequency of stimulation to isolate each photoactivation /PSP instance, including a post-response baseline period (where cell was at resting). To determine sensitivity for a certain number of trials averaged (N_t_), the z-score of the response amplitude (or response SNR) for an N_t_-averaged trace was calculated relative to the distribution of bootstrap-sampled (n = 500) averages of the baseline (no-response) period in the averaged trace. This comparison was performed 50 times for N_t_ randomly sampled PSPs in the photoactivation series, and the mean of the 50 resulting SNR measurements was used to determine sensitivity.

#### Evaluation of optimal number of PSP trials to average

Because some cells in our MOSAIX experiments showed non-stationarity in PSP amplitude over the course of the photoactivation series, we first determined the optimal number of consecutive PSPs to average (N_o_; see Supplementary Text) before generating the trial-averaged PSP for each cell. Specifically, we computed the z-score of the response amplitude SNR for traces averaged over N_t_ consecutive trials, relative to the mean and SEM of the baseline period (recorded while the cell was at rest). N_o_ was defined as the value of N_t_ that yielded the largest-magnitude SNR response.

To quantify the false-positive rate associated with a given SNR threshold for classifying cells as “responding” or “not responding”, we additionally performed bootstrap resampling (n = 500) from the baseline period of the N_o_-averaged trace to generate a null distribution of SNR values. A threshold of SNR z ≤ −3.75 yielded a false-positive rate of < 0.07 across all imaged cells. Accordingly, a cell was classified as “responding” only if the SNR of its N_o_-averaged response met or exceeded this threshold (i.e., SNR ≤ −3.75).

#### Multi-channel HCR image unmixing

In each round of HCR, 5 genes were probed across 5 fluorescent channels: AlexaFluor 488 (AF488), AF514, AF546, AF594, and AF647. Genes imaged in spectrally adjacent channels were always chosen to be non-overlapping in expression. Signal from AF488 was detectable in the imaged AF514 channel due to overlapping excitation and emission spectra for these fluorophores. This was similarly the case for AF594 signal in the imaged AF546 channel. To spectrally unmix these signals, the signal from the source channel was scaled down (my multiplying all pixel values by a fraction scalar, such as 0.7) and subtracted from the recipient channel. The scalar value used was incrementally adjusted until the signal subtracted from the recipient image yielded pixel values approximating the tissue autofluorescence intensity in that channel. Lipofuscin signal was also removed from all channel images by evaluating the 5-channel spectral profile of pixels containing lipofuscin signal and replacing all pixels matching this profile with pixel intensities that matched the tissue autofluorescence for the corresponding channel.

#### Multi-round HCR image registration

Images of NeuroTrace 435/455 or Snap25 mFISH signal were used for registering pre-HCR confocal images and HCR mFISH images across imaging rounds. Raw images were first preprocessed to enhance local contrast and remove background signal. Local intensity normalization was applied using a block-based contrast stretching algorithm, followed by background subtraction using a rolling ball algorithm with a 40-pixel radius. Prior to automated registration, an initial affine transformation was estimated from manually selected corresponding landmarks in the fixed and moving images and used to compute an initial affine transform. This transform provided a coarse alignment to facilitate subsequent intensity-based gradient descent. These corresponding points were further used as a metric in the automated gradient descent. Image registration was performed using the elastix software package^60^ and was comprised of multiple steps: 1) Affine refinement: an affine transform was optimized combining mutual information between intensities and a corresponding-points Euclidean distance metric. 2) Nonrigid B-spline registration with penalties: a penalized B-spline deformation was applied using 4 resolutions depending on the dataset, optimizing mutual information alongside bending energy and point correspondence penalties. 3) Final nonrigid B-spline refinement: a final non-penalized B-spline registration stage was performed to fine-tune alignment using mutual information alone at the final B-spline resolution.

#### mFISH image segmentation

To create individual cell ROIs in initial HCR mFISH runs, the AlexaFluor 488 (AF488) HCR channel containing signal from *Snap25* transcripts was used for segmentation. For MOSAIX samples, the AF514 HCR channel containing signal for *Vip* transcripts and spectral bleed from AF488 Snap25 signal was used for segmentation because a subset of *Vip*+ cells did not show *Snap25* expression. The *Vip*+*Snap25* image was first background-subtracted in FIJI^59^ using a rolling ball algorithm with 50-100 px radius. Sample images were hand annotated to mark cell boundaries and the resulting ROIs were used to train a segmentation model in Cellpose2 or Cellpose3^61,62^. Confocal volume images were then segmented with z-axis segmentation performed using the intersection-over-union evaluation with a 70% threshold. Flow threshold and cell probability thresholds were kept at default values (0.4, 0.0 respectively), as were lower and upper pixel intensity percentiles (1.0, 99.0 respectively). The pixel value assigned to each cell’s ROI was used as the cell ID for all downstream voltage imaging and mFISH classification analyses.

#### Binary annotation of mFISH data and classification

In HCR mFISH experiments used evaluate cell type, segmented cell ROIs from full HCR mFISH volumes were annotated in a binary fashion for each gene, such that each cell received a “positive” or “negative” label for each marker gene that was probed. For experiments where all cells in a tissue sample were classified, annotation was performed using the “Object Classification” module in Ilastik^63^, which allowed the user to select examples of “positive” and “negative” cells in the sample to train a random forest classifier that evaluated all available features in the module. This classifier was then applied to annotate the remaining population of segmented neurons in the tissue volume, and could be exported to classify cells across all experiments. In MOSAIX experiments, binary annotation of gene expression was performed on a per-cell and per-gene basis. Briefly, using a custom GUI, the full image volume containing a cell’s HCR mFISH signal was evaluated against the shape of the segmentation ROI and against the Voltron2-JF585 signal (to avoid incorrectly annotating immediately adjacent cells). For a given gene, mFISH signal in each cell ROI was first quantitatively evaluated using the ratio of the mean intensity of mFISH signal inside the ROI vs. in non-ROI pixels found inside a 100×100 pixel area immediately surrounding the cell’s ROI. This allowed all cells to be rank-ordered by expression “intensity”, and an initial binary expression threshold to be set. All cells above and below this threshold were then manually evaluated to ensure a “positive” or “negative” label was appropriately assigned. A binary annotation matrix for HCR cells (structured as cells x genes) was generated from either Ilastik or manual annotation data. To create comparable binary expression patterns in ABC Atlas scRNA-seq data^11^, we generated a population of “meta cells”: cells from the scRNA-seq dataset – specifically the “WMB-10Xv2-Isocortex-1” dataset – whose normalized counts values were summed with those of their 9 nearest neighbors in PCA space (30 PCs, identified using Seurat^64^). In this way, each meta cell represented 10 cells in the scRNA-seq dataset. This population of meta cells showed clear multimodal distributions of normalized counts for each gene, composed of low-and high-expressing cells. To draw binarization boundaries between these portions of the distributions, we fit a gaussian mixture model to the distributions (excluding the peak representing zero counts), using the minimum number of gaussians above which decreasing Akaike and Bayes information criteria (AIC/BIC) scores leveled off. We then determined the intersection points between fit gaussians and used these points as thresholds for binarization. For distributions with more than two fit gaussians, the intersection point used was selected such that the data would be divided approximately in half by the threshold. Cells with normalized counts values above the intersection point were considered “positive” for a gene, with cells below labeled as “negative”. For each cell type, this allowed us to determine the proportion of cells of each type that express each of the 17 genes probed by HCR – essentially determining how likely a given cell type is to express a particular gene. These proportions were used as probabilities in a Bernoulli Bayes Classifier to classify each cell imaged in HCR experiments into a taxonomic cell type. The evaluated cells’ laminar depths were used to select depth-specific cell type probabilities (used as the prior probability) derived from ABC MERFISH data, with each cell type having a baseline probability of 0.2% at each depth. The classifier evaluated classification probabilities as follows:

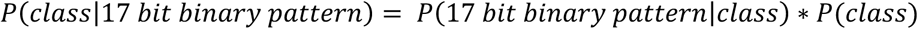

The classification was performed in a hierarchical fashion, such that the first evaluation was for Glut vs. GABA neurotransmitter type. If a cell was classified as Glut, it would go on to be subclassed into types forming the Glut class. At each evaluation node, the cell type that received the highest probability score became a cell’s assigned type. Excitatory neurons were classed down to the level of “supertype”, while inhibitory neurons were classed down the “subclass” level.

#### Columnar CheRiff signal quantification

A maximum intensity projection image of CheRiff-EGFP signal was generated across the z-axis of a confocal CheRiff image. The x-y location of each Voltron2-positive cell from the same confocal volume was then mapped onto this 2D image. A 75 µm-wide column perpendicular to the pial surface and spanning from pia to the corpus callosum, was centered in x around the cell of interest. The mean intensity of CheRiff signal in this column was measured, and a normalized intensity value was generated by normalizing each column’s mean to the mean of the brightest column in the same slice. In MOSAIX data, response amplitudes were normalized to columnar CheRiff intensity by dividing median-normalized response amplitudes (response amplitudes divided by the median amplitude of responding cells in the same slice) by the normalized columnar CheRiff value.

#### mFISH and voltage imaging data registration

To register widefield 2D images of Voltron2-JF585 signal to 3D confocal volumes of the same signal, the general registration pipeline described for multi-round HCR mFISH image registration was used, with modifications to account for dimensionality. Confocal image stacks were first oriented based on manually selected landmarks and z-projected based on each z-plane correlation with the widefield image. The confocal stacks were additionally Gaussian blurred before running elastix to calculate the final B-spline transform. Finally, Voltron2+ segmentation masks generated from mFISH confocal data were registered to Voltron2JF585 widefield images using this same transform, and masks were sub-selected based on alignment to distinguishable cells in the widefield image. These aligned masks were used to extract timeseries data for Voltron2-JF585 in MOSAIX experiments.

### QUANTIFICATION AND STATISTICAL ANALYSIS

#### Voltage imaging sensitivity evaluation

Both whole-cell recording and voltage imaging were segmented in time based on the frequency of stimulation to isolate each photoactivation /PSP instance, including a post-response baseline period (where cell was at resting). The z-score of the response amplitude (or response SNR) for an N-trial-averaged trace was calculated relative to the distribution of bootstrap-sampled (n = 500) averages of the baseline (no-response) period in the averaged trace. This comparison was performed 50 times for randomly sampled N-trial PSPs in the photoactivation series, and the mean of the 50 resulting SNR measurements was used to determine sensitivity.

#### Evaluation of optimal number of PSP trials to average

We computed the z-score of the response amplitude SNR for traces averaged over consecutive numbers of trials, relative to the mean and SEM of the baseline (no-reponse) period. Cells were subsequently classified as “responding” if the SNR of its averaged response met or exceeded z = –3.75 relative to a null distribution of SNR values generated from the bootstrap sampled (n = 500) no-response period.

#### Comparisons of cell type response amplitudes

Differences in PSP amplitudes between two cell types were assessed with two-tailed Student’s t-tests. For comparisons involving three cell types, we used one-way ANOVA followed by post hoc pairwise tests.

#### Gaussian mixture model parameters

To model the distribution of “meta cell” counts (excluding the component corresponding to zero counts), we fitted Gaussian mixture models with varying numbers of components and compared models using the Akaike Information Criterion (AIC) and the Bayesian Information Criterion (BIC). We selected the smallest number of Gaussian components beyond which AIC and BIC showed negligible improvement (i.e., where the score curves asymptoted or “elbowed”), preferring the simpler model when criteria were similar.

#### Reporting of statistical parameters

All replicate numbers (n) and statistical parameters—standard error of the mean (SEM), standard deviation (SD), interquartile range (IQR), and confidence intervals (CI)—are reported in the main text, figure legends, and in the figures.

#### Bayesian classification of cell types in mFISH data

The proportions of “meta cell” types that expressed each mFISH marker gene were used as probabilities in a Bernoulli Bayes Classifier to classify each cell imaged in HCR experiments into a taxonomic cell type. The evaluated cells’ laminar depths were used to select depth-specific cell type probabilities (used as the prior probability), with each cell type having a baseline probability of 0.2% at each depth. The classifier evaluated classification probabilities as follows:

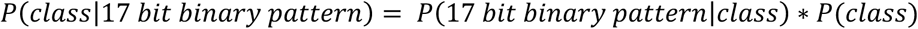

## KEY RESOURCES TABLE

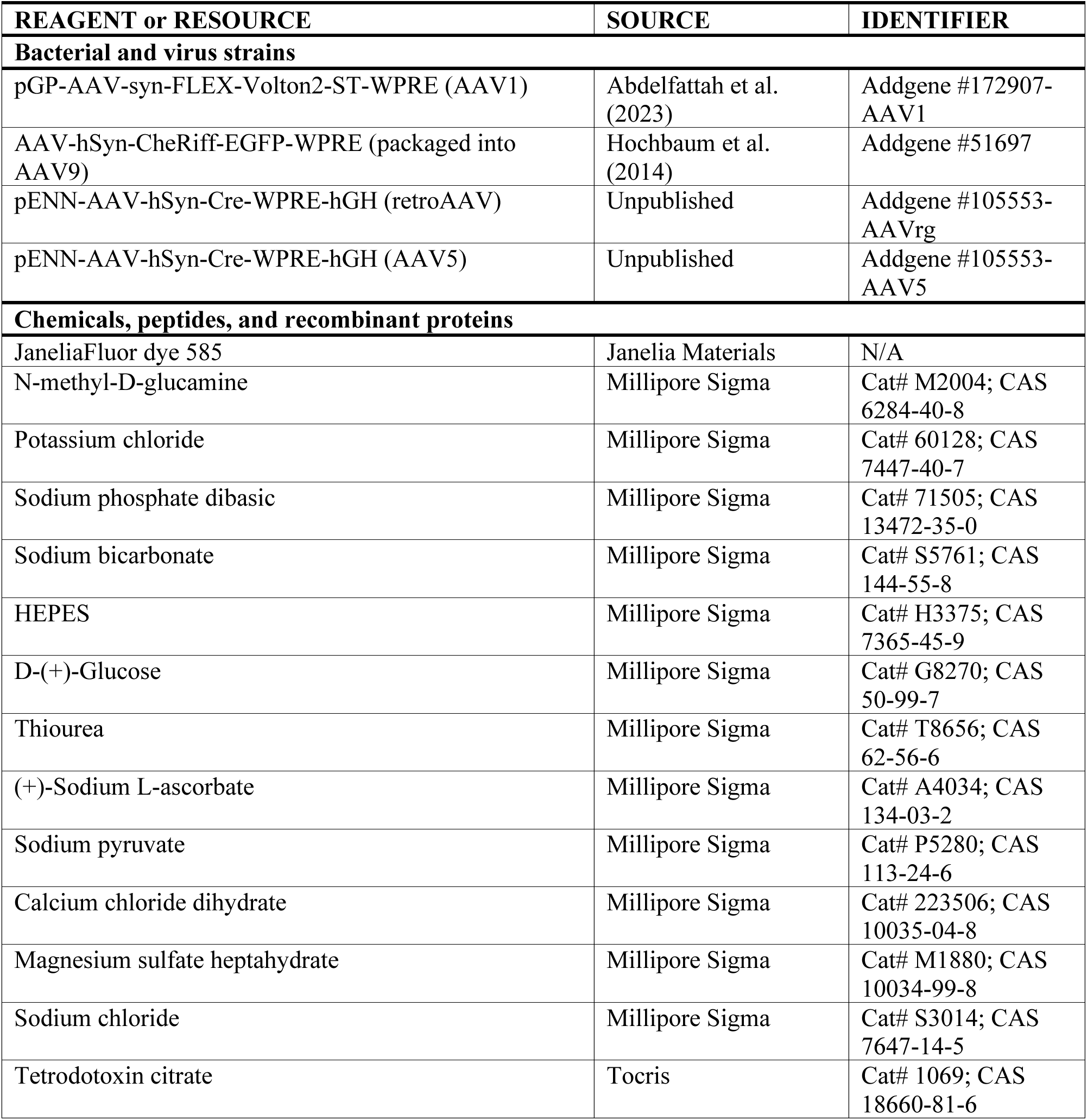
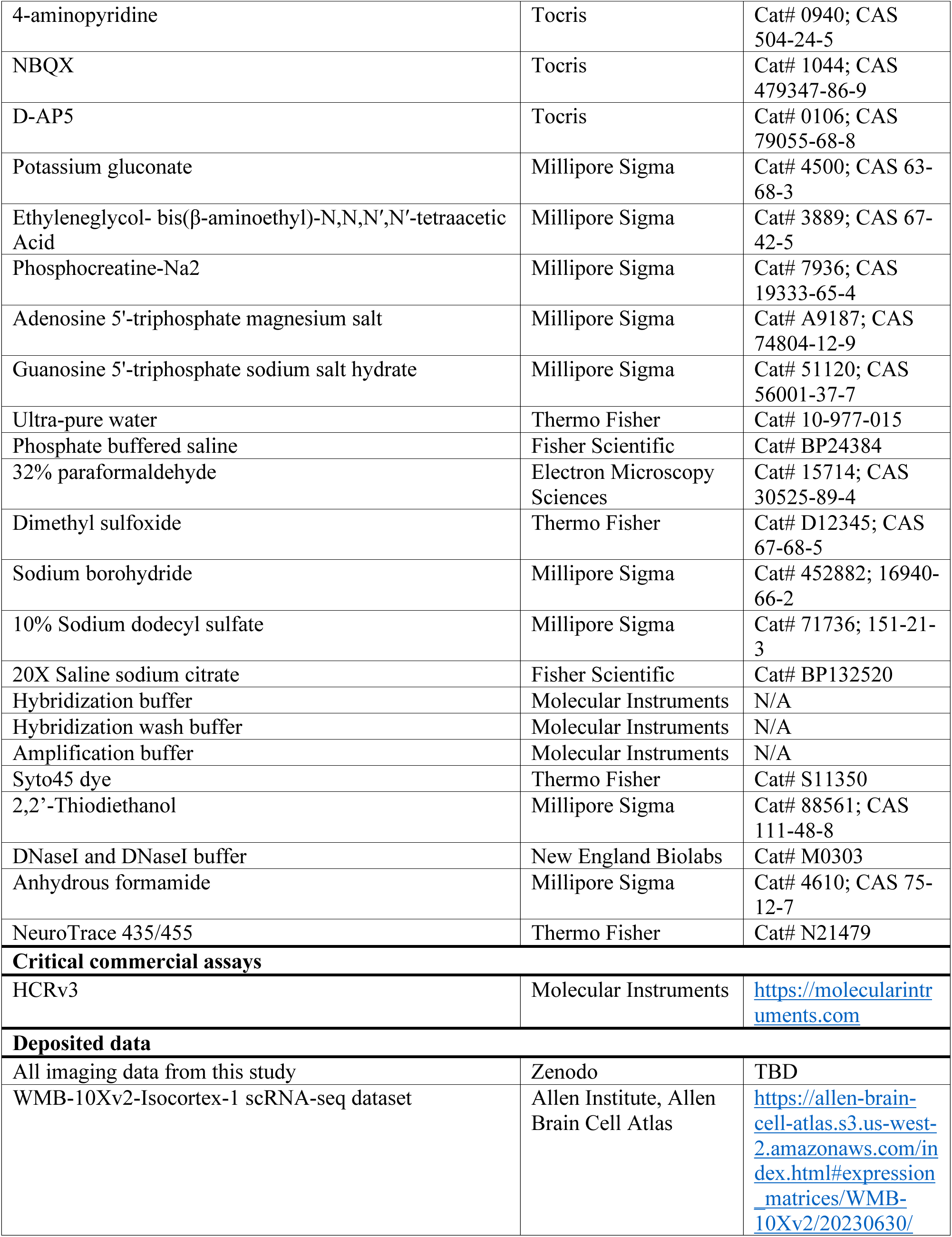
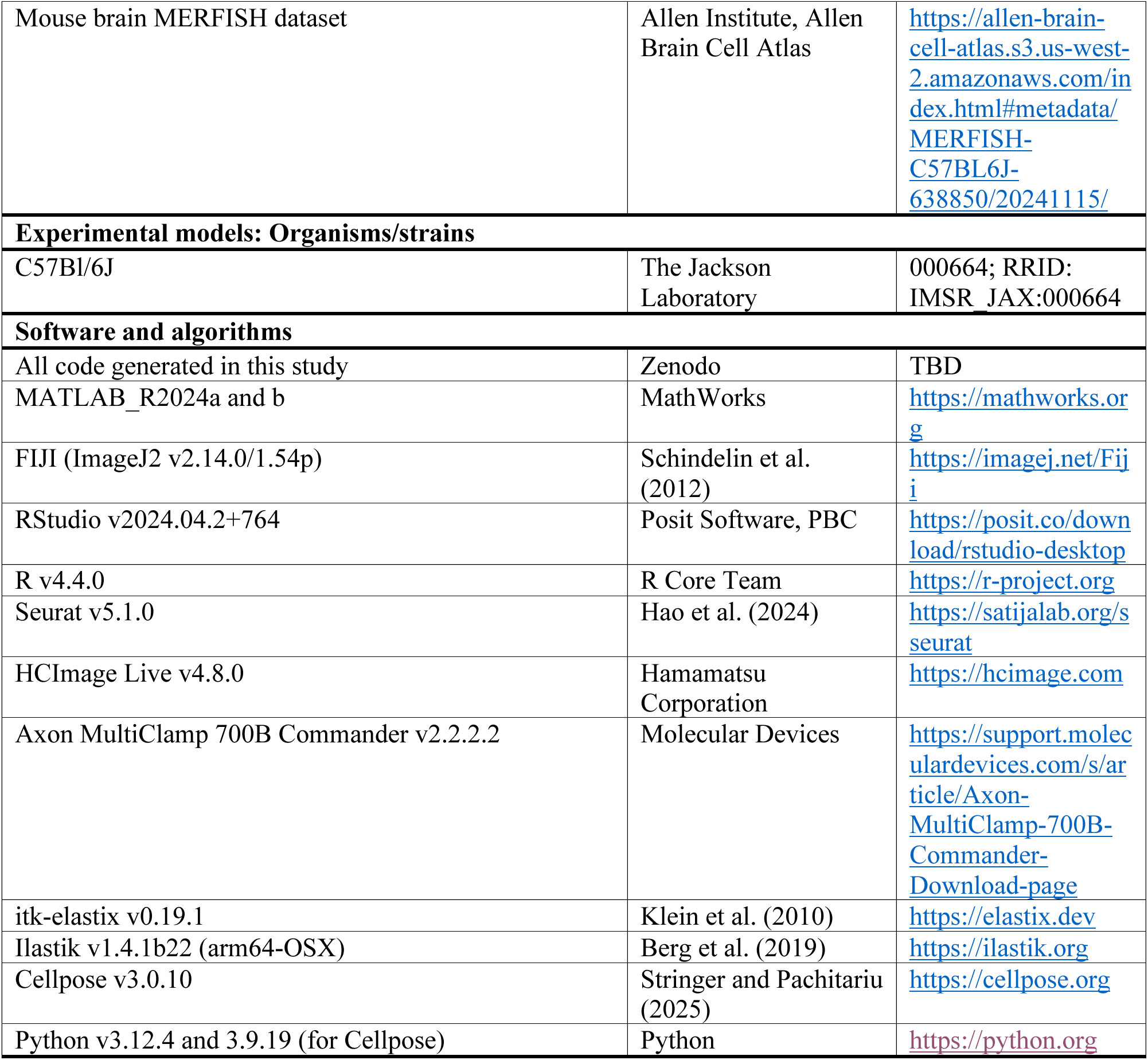

## Notes

### Competing Interest Statement

The authors have declared no competing interest.

### Summary of Updates

Clearer articulation of novelty. We have revised the manuscript to better highlight the innovations of MOSAIX (renamed from OCM-FISH). These include: (1) the demonstration of reliable, parallelized detection of sub-millivolt postsynaptic potentials with voltage imaging; (2) an accessible mFISH pipeline for identifying cell types in thick tissue; and (3) a novel, scalable pipeline approach for integrating connectivity measurements with cell-type identification. Enhanced reproducibility and rigor. We now include comprehensive analyses showing that cell-type-specific connectivity patterns are highly reproducible across MOSAIX experiments.

## References

1. Marr, D. (1969). A theory of cerebellar cortex. J. Physiol. 202, 437–470. 10.1113/jphysiol.1969.sp008820.

2. Lorente De Nó, R. (1934). Studies on the structure of the cerebral cortex. II. Continuation of the study of the ammonic system. J. Für Psychol. Neurol. 46, 113–177.

3. Green, J.D., and Adey, W.R. (1956). Electrophysiological studies of hippocampal connections and excitability. Electroencephalogr. Clin. Neurophysiol. 8, 245–262. 10.1016/0013-4694(56)90117-1.

4. Albin, R.L., Young, A.B., and Penney, J.B. (1989). The functional anatomy of basal ganglia disorders. Trends Neurosci. 12, 366–375. 10.1016/0166-2236(89)90074-X.

5. DeLong, M.R. (1990). Primate models of movement disorders of basal ganglia origin. Trends Neurosci. 13, 281–285. 10.1016/0166-2236(90)90110-V.

6. Hubel, D.H., and Wiesel, T.N. (1962). Receptive fields, binocular interaction and functional architecture in the cat’s visual cortex. J. Physiol. 160, 106. 10.1113/jphysiol.1962.sp006837.

7. Hubel, D.H., and Wiesel, T.N. (1968). Receptive fields and functional architecture of monkey striate cortex. J. Physiol. 195, 215–243. 10.1113/jphysiol.1968.sp008455.

8. Mountcastle, V.B. (1997). The columnar organization of the neocortex. Brain 120, 701–722. 10.1093/brain/120.4.701.

9. Famiglietti, E.V., and Kolb, H. (1976). Structural Basis for ON-and OFF-Center Responses in Retinal Ganglion Cells. Science 194, 193–195. 10.1126/science.959847.

10. Yao, Z., van Velthoven, C.T.J., Nguyen, T.N., Goldy, J., Sedeno-Cortes, A.E., Baftizadeh, F., Bertagnolli, D., Casper, T., Chiang, M., Crichton, K., et al. (2021). A taxonomy of transcriptomic cell types across the isocortex and hippocampal formation. Cell 184, 3222–3241.e26. 10.1016/j.cell.2021.04.021.

11. Yao, Z., van Velthoven, C.T.J., Kunst, M., Zhang, M., McMillen, D., Lee, C., Jung, W., Goldy, J., Abdelhak, A., Aitken, M., et al. (2023). A high-resolution transcriptomic and spatial atlas of cell types in the whole mouse brain. Nature 624, 317–332. 10.1038/s41586-023-06812-z.

12. Perin, R., Berger, T.K., and Markram, H. (2011). A synaptic organizing principle for cortical neuronal groups. Proc. Natl. Acad. Sci. 108, 5419–5424. 10.1073/pnas.1016051108.

13. Jiang, X., Shen, S., Cadwell, C.R., Berens, P., Sinz, F., Ecker, A.S., Patel, S., and Tolias, A.S. (2015). Principles of connectivity among morphologically defined cell types in adult neocortex. Science 350, aac9462. 10.1126/science.aac9462.

14. Campagnola, L., Seeman, S.C., Chartrand, T., Kim, L., Hoggarth, A., Gamlin, C., Ito, S., Trinh, J., Davoudian, P., Radaelli, C., et al. (2022). Local connectivity and synaptic dynamics in mouse and human neocortex. Science 375, eabj5861. 10.1126/science.abj5861.

15. Petreanu, L., Huber, D., Sobczyk, A., and Svoboda, K. (2007). Channelrhodopsin-2–assisted circuit mapping of long-range callosal projections. Nat. Neurosci. 10, 663–668. 10.1038/nn1891.

16. Petreanu, L., Mao, T., Sternson, S.M., and Svoboda, K. (2009). The subcellular organization of neocortical excitatory connections. Nature 457, 1142–1145. 10.1038/nature07709.

17. Fan, L.Z., Nehme, R., Adam, Y., Jung, E.S., Wu, H., Eggan, K., Arnold, D.B., and Cohen, A.E. (2018). All-optical synaptic electrophysiology probes mechanism of ketamine-induced disinhibition. Nat. Methods 15, 823–831. 10.1038/s41592-018-0142-8.

18. Abdelfattah, A.S., Zheng, J., Singh, A., Huang, Y.-C., Reep, D., Tsegaye, G., Tsang, A., Arthur, B.J., Rehorova, M., Olson, C.V.L., et al. (2023). Sensitivity optimization of a rhodopsin-based fluorescent voltage indicator. Neuron 111, 1547–1563.e9. 10.1016/j.neuron.2023.03.009.

19. Hao, Y.A., Lee, S., Roth, R.H., Natale, S., Gomez, L., Taxidis, J., O’Neill, P.S., Villette, V., Bradley, J., Wang, Z., et al. (2024). A fast and responsive voltage indicator with enhanced sensitivity for unitary synaptic events. Neuron 112, 3680–3696.e8. 10.1016/j.neuron.2024.08.019.

20. Liu, Z., Lu, X., Villette, V., Gou, Y., Colbert, K.L., Lai, S., Guan, S., Land, M.A., Lee, J., Assefa, T., et al. (2022). Sustained deep-tissue voltage recording using a fast indicator evolved for two-photon microscopy. Cell 185, 3408–3425.e29. 10.1016/j.cell.2022.07.013.

21. Gong, Y., Huang, C., Li, J.Z., Grewe, B.F., Zhang, Y., Eismann, S., and Schnitzer, M.J. (2015). High-speed recording of neural spikes in awake mice and flies with a fluorescent voltage sensor. Science 350, 1361–1366. 10.1126/science.aab0810.

22. Adam, Y., Kim, J.J., Lou, S., Zhao, Y., Xie, M.E., Brinks, D., Wu, H., Mostajo-Radji, M.A., Kheifets, S., Parot, V., et al. (2019). Voltage imaging and optogenetics reveal behaviour-dependent changes in hippocampal dynamics. Nature 569, 413–417. 10.1038/s41586-019-1166-7.

23. Piatkevich, K.D., Bensussen, S., Tseng, H., Shroff, S.N., Lopez-Huerta, V.G., Park, D., Jung, E.E., Shemesh, O.A., Straub, C., Gritton, H.J., et al. (2019). Population imaging of neural activity in awake behaving mice. Nature 574, 413–417. 10.1038/s41586-019-1641-1.

24. Fan, L.Z., Kheifets, S., Böhm, U.L., Wu, H., Piatkevich, K.D., Xie, M.E., Parot, V., Ha, Y., Evans, K.E., Boyden, E.S., et al. (2020). All-Optical Electrophysiology Reveals the Role of Lateral Inhibition in Sensory Processing in Cortical Layer 1. Cell 180, 521–535.e18. 10.1016/j.cell.2020.01.001.

25. Kim, T.H., and Schnitzer, M.J. (2022). Fluorescence imaging of large-scale neural ensemble dynamics. Cell 185, 9–41. 10.1016/j.cell.2021.12.007.

26. Abdelfattah, A.S., Zheng, J., Singh, A., Huang, Y.-C., Reep, D., Tsegaye, G., Tsang, A., Arthur, B.J., Rehorova, M., Olson, C.V.L., et al. (2023). Sensitivity optimization of a rhodopsin-based fluorescent voltage indicator. Neuron 111, 1547–1563.e9. 10.1016/j.neuron.2023.03.009.

27. Hao, Y.A., Lee, S., Roth, R.H., Natale, S., Gomez, L., Taxidis, J., O’Neill, P.S., Villette, V., Bradley, J., Wang, Z., et al. (2024). A fast and responsive voltage indicator with enhanced sensitivity for unitary synaptic events. Neuron 112, 3680–3696.e8. 10.1016/j.neuron.2024.08.019.

28. Hochbaum, D.R., Zhao, Y., Farhi, S.L., Klapoetke, N., Werley, C.A., Kapoor, V., Zou, P., Kralj, J.M., Maclaurin, D., Smedemark-Margulies, N., et al. (2014). All-optical electrophysiology in mammalian neurons using engineered microbial rhodopsins. Nat. Methods 11, 825–833. 10.1038/nmeth.3000.

29. Grimm, J.B., Tkachuk, A.N., Xie, L., Choi, H., Mohar, B., Falco, N., Schaefer, K., Patel, R., Zheng, Q., Liu, Z., et al. (2020). A general method to optimize and functionalize red-shifted rhodamine dyes. Nat. Methods 17, 815–821. 10.1038/s41592-020-0909-6.

30. Cruikshank, S.J., Urabe, H., Nurmikko, A.V., and Connors, B.W. (2010). Pathway-specific feedforward circuits between thalamus and neocortex revealed by selective optical stimulation of axons. Neuron 65, 230–245. 10.1016/j.neuron.2009.12.025.

31. Guo, K., Yamawaki, N., Svoboda, K., and Shepherd, G.M.G. (2018). Anterolateral Motor Cortex Connects with a Medial Subdivision of Ventromedial Thalamus through Cell Type-Specific Circuits, Forming an Excitatory Thalamo-Cortico-Thalamic Loop via Layer 1 Apical Tuft Dendrites of Layer 5B Pyramidal Tract Type Neurons. J. Neurosci. 38, 8787–8797. 10.1523/JNEUROSCI.1333-18.2018.

32. Yang, W., Tipparaju, S.L., Chen, G., and Li, N. (2022). Thalamus-driven functional populations in frontal cortex support decision-making. Nat. Neurosci. 25, 1339–1352. 10.1038/s41593-022-01171-w.

33. Shu, Y., Yu, Y., Yang, J., and McCormick, D.A. (2007). Selective control of cortical axonal spikes by a slowly inactivating K+ current. Proc. Natl. Acad. Sci. 104, 11453–11458. 10.1073/pnas.0702041104.

34. Chen, K.H., Boettiger, A.N., Moffitt, J.R., Wang, S., and Zhuang, X. (2015). Spatially resolved, highly multiplexed RNA profiling in single cells. Science 348, aaa6090. 10.1126/science.aaa6090.

35. Shah, S., Lubeck, E., Zhou, W., and Cai, L. (2016). In Situ Transcription Profiling of Single Cells Reveals Spatial Organization of Cells in the Mouse Hippocampus. Neuron 92, 342–357. 10.1016/j.neuron.2016.10.001.

36. Lee, J.H., Daugharthy, E.R., Scheiman, J., Kalhor, R., Ferrante, T.C., Terry, R., Turczyk, B.M., Yang, J.L., Lee, H.S., Aach, J., et al. (2015). Fluorescent in situ sequencing (FISSEQ) of RNA for gene expression profiling in intact cells and tissues. Nat. Protoc. 10, 442–458. 10.1038/nprot.2014.191.

37. Wang, Y., Eddison, M., Fleishman, G., Weigert, M., Xu, S., Wang, T., Rokicki, K., Goina, C., Henry, F.E., Lemire, A.L., et al. (2021). EASI-FISH for thick tissue defines lateral hypothalamus spatio-molecular organization. Cell 184, 6361–6377.e24. 10.1016/j.cell.2021.11.024.

38. Choi, H.M.T., Schwarzkopf, M., Fornace, M.E., Acharya, A., Artavanis, G., Stegmaier, J., Cunha, A., and Pierce, N.A. (2018). Third-generation in situ hybridization chain reaction: multiplexed, quantitative, sensitive, versatile, robust. Development 145. 10.1242/dev.165753.

39. Svensson, V., Natarajan, K.N., Ly, L.-H., Miragaia, R.J., Labalette, C., Macaulay, I.C., Cvejic, A., and Teichmann, S.A. (2017). Power analysis of single-cell RNA-sequencing experiments. Nat. Methods 14, 381–387. 10.1038/nmeth.4220.

40. Baran-Gale, J., Chandra, T., and Kirschner, K. (2018). Experimental design for single-cell RNA sequencing. Brief. Funct. Genomics 17, 233–239. 10.1093/bfgp/elx035.

41. Wang, X., He, Y., Zhang, Q., Ren, X., and Zhang, Z. (2021). Direct Comparative Analyses of 10X Genomics Chromium and Smart-seq2. Genomics Proteomics Bioinformatics 19, 253–266. 10.1016/j.gpb.2020.02.005.

42. Hooks, B.M., Mao, T., Gutnisky, D.A., Yamawaki, N., Svoboda, K., and Shepherd, G.M.G. (2013). Organization of Cortical and Thalamic Input to Pyramidal Neurons in Mouse Motor Cortex. J. Neurosci. 33, 748–760. 10.1523/JNEUROSCI.4338-12.2013.

43. Yamawaki, N., Borges, K., Suter, B.A., Harris, K.D., and Shepherd, G.M.G. (2014). A genuine layer 4 in motor cortex with prototypical synaptic circuit connectivity. eLife 3, e05422. 10.7554/eLife.05422.

44. Muñoz-Castañeda, R., Zingg, B., Matho, K.S., Chen, X., Wang, Q., Foster, N.N., Li, A., Narasimhan, A., Hirokawa, K.E., Huo, B., et al. (2021). Cellular anatomy of the mouse primary motor cortex. Nature 598, 159–166. 10.1038/s41586-021-03970-w.

45. Economo, M.N., Viswanathan, S., Tasic, B., Bas, E., Winnubst, J., Menon, V., Graybuck, L.T., Nguyen, T.N., Smith, K.A., Yao, Z., et al. (2018). Distinct descending motor cortex pathways and their roles in movement. Nature 563, 79–84. 10.1038/s41586-018-0642-9.

46. Yamawaki, N., and Shepherd, G.M.G. (2015). Synaptic circuit organization of motor corticothalamic neurons. J. Neurosci. Off. J. Soc. Neurosci. 35. 10.1523/JNEUROSCI.4023-14.2015.

47. Chou, C.Y.C., Wong, H.H.W., Guo, C., Boukoulou, K.E., Huang, C., Jannat, J., Klimenko, T., Li, V.Y., Liang, T.A., Wu, V.C., et al. (2025). Principles of visual cortex excitatory microcircuit organization. The Innovation 6, 100735. 10.1016/j.xinn.2024.100735.

48. Csillag, V., Bizzozzero, M.H., Noble, J.C., Reinius, B., and Fuzik, J. (2023). Voltage-Seq: all-optical postsynaptic connectome-guided single-cell transcriptomics. Nat. Methods 20, 1409–1416. 10.1038/s41592-023-01965-1.

49. Weber, T.D., Moya, M.V., Kılıç, K., Mertz, J., and Economo, M.N. (2023). High-speed multiplane confocal microscopy for voltage imaging in densely labeled neuronal populations. Nat. Neurosci. 26, 1642–1650. 10.1038/s41593-023-01408-2.

50. Xiao, S., Cunningham, W.J., Kondabolu, K., Lowet, E., Moya, M.V., Mount, R.A., Ravasio, C., Bortz, E., Shaw, D., Economo, M.N., et al. (2024). Large-scale deep tissue voltage imaging with targeted-illumination confocal microscopy. Nat. Methods 21, 1094–1102. 10.1038/s41592-024-02275-w.

51. Salter, E.W., Galeazzi, F., Sheinberg, D., Test, M., Amin, K., Paradiso, B., and Abdelfattah, A.S. (2024). Development of a red-shifted chemigenetic voltage indicator.

52. Markram, H., Lübke, J., Frotscher, M., Roth, A., and Sakmann, B. (1997). Physiology and anatomy of synaptic connections between thick tufted pyramidal neurones in the developing rat neocortex. J. Physiol. 500, 409–440. 10.1113/jphysiol.1997.sp022031.

53. Song, S., Sjöström, P.J., Reigl, M., Nelson, S., and Chklovskii, D.B. (2005). Highly Nonrandom Features of Synaptic Connectivity in Local Cortical Circuits. PLOS Biol. 3, e68. 10.1371/journal.pbio.0030068.

54. Jiang, X., Shen, S., Cadwell, C.R., Berens, P., Sinz, F., Ecker, A.S., Patel, S., and Tolias, A.S. (2015). Principles of connectivity among morphologically defined cell types in adult neocortex. Science 350, aac9462. 10.1126/science.aac9462.

55. Fang, R., Halpern, A.R., Rahman, M.M., Huang, Z., Lei, Z., Hell, S.J., Dulac, C., and Zhuang, X. (2024). Three-dimensional single-cell transcriptome imaging of thick tissues. eLife 12. 10.7554/eLife.90029.2.

56. Ting, J.T., Lee, B.R., Chong, P., Soler-Llavina, G., Cobbs, C., Koch, C., Zeng, H., and Lein, E. (2018). Preparation of Acute Brain Slices Using an Optimized N-Methyl-D-glucamine Protective Recovery Method. JoVE J. Vis. Exp., e53825. 10.3791/53825.

57. Lubeck, E., Coskun, A.F., Zhiyentayev, T., Ahmad, M., and Cai, L. (2014). Single-cell in situ RNA profiling by sequential hybridization. Nat. Methods 11, 360–361. 10.1038/nmeth.2892.

58. Pnevmatikakis, E.A., and Giovannucci, A. (2017). NoRMCorre: An online algorithm for piecewise rigid motion correction of calcium imaging data. J. Neurosci. Methods 291, 83–94. 10.1016/j.jneumeth.2017.07.031.

59. Schindelin, J., Arganda-Carreras, I., Frise, E., Kaynig, V., Longair, M., Pietzsch, T., Preibisch, S., Rueden, C., Saalfeld, S., Schmid, B., et al. (2012). Fiji: an open-source platform for biological-image analysis. Nat. Methods 9, 676–682. 10.1038/nmeth.2019.

60. Klein, S., Staring, M., Murphy, K., Viergever, M.A., and Pluim, J.P.W. (2010). elastix: A Toolbox for Intensity-Based Medical Image Registration. IEEE Trans. Med. Imaging 29, 196–205. 10.1109/TMI.2009.2035616.

61. Pachitariu, M., and Stringer, C. (2022). Cellpose 2.0: how to train your own model. Nat. Methods 19, 1634–1641. 10.1038/s41592-022-01663-4.

62. Stringer, C., and Pachitariu, M. (2025). Cellpose3: one-click image restoration for improved cellular segmentation. Nat. Methods 22, 592–599. 10.1038/s41592-025-02595-5.

63. Berg, S., Kutra, D., Kroeger, T., Straehle, C.N., Kausler, B.X., Haubold, C., Schiegg, M., Ales, J., Beier, T., Rudy, M., et al. (2019). ilastik: interactive machine learning for (bio)image analysis. Nat. Methods 16, 1226–1232. 10.1038/s41592-019-0582-9.

64. Hao, Y., Stuart, T., Kowalski, M.H., Choudhary, S., Hoffman, P., Hartman, A., Srivastava, A., Molla, G., Madad, S., Fernandez-Granda, C., et al. (2024). Dictionary learning for integrative, multimodal and scalable single-cell analysis. Nat. Biotechnol. 42, 293–304. 10.1038/s41587-023-01767-y.

