## Supplementary Text and Figures for "A scalable, all-optical method for mapping synaptic connectivity with cell-type specificity"

### Pharmacological blockers cause attenuation of PSPs in repetitive photostimulation experiments.

In optical circuit mapping experiments, postsynaptic responses to repeated 2 Hz presynaptic axon photoactivation showed consistent amplitudes in the absence of pharmacological blockers (**Figure 5a,b**), but we observed modest attenuation of postsynaptic potential (PSP) amplitude after repetitive photoactivation of motor thalamus (MThal) axons in the presence of 4-AP in some neurons, which was accelerated by the subsequent application of tetrodotoxin (TTX). Increasing concentrations of TTX showed increasing rates of attenuation, but 200 nM TTX appeared to provide a stable PSP amplitude in the second half of stimulation series (**Figure S5b**). For this reason, we opted to use 200 nM TTX for most circuit mapping experiments, except in cases where this concentration resulted in inhibitory responses that were likely poly-synaptic (**Figure S5c**), in which case a higher concentration was titrated into the extracellular solution until these inhibitory responses were largely abolished. The biophysical mechanism resulting from attenuation in the presence of 4-aminopyridine and TTX, but not otherwise, is not clear.

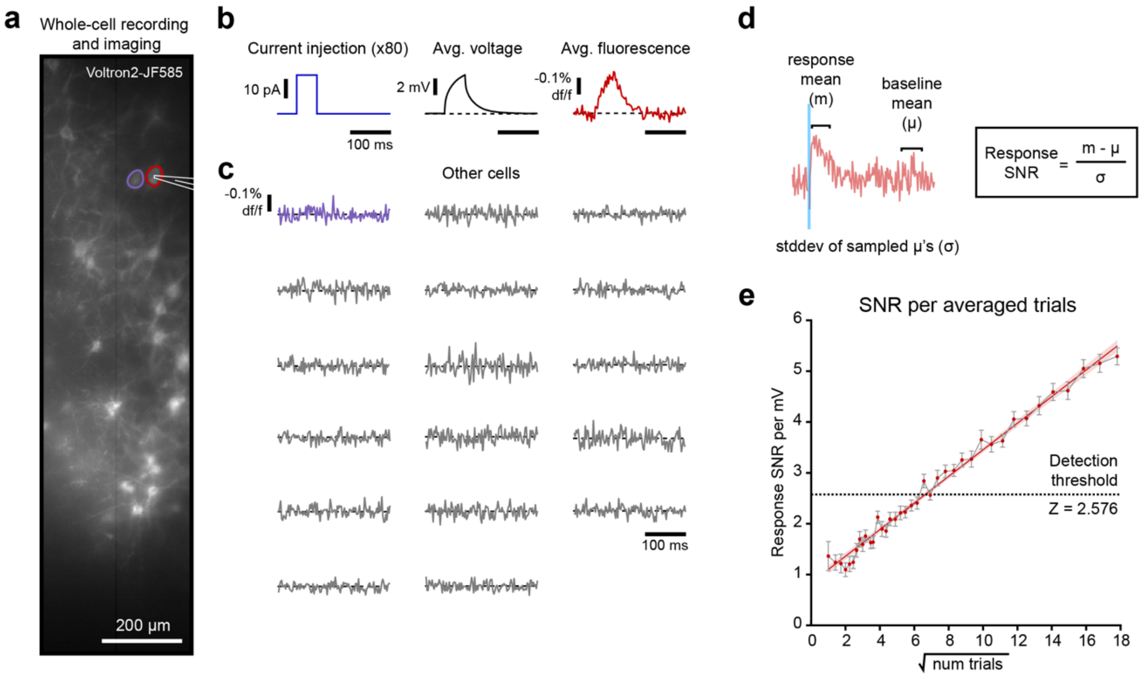


*Figure S1 – Whole-cell recording with voltage imaging allows for sensitivity measurements.* (***a***) Widefield image of Voltron2-JF585+ cells with two cells highlighted, including a cell that was patched for simultaneous whole-cell recording. (***b***) Short current injections (blue) were performed in patched cells to test for efficacious patch in both membrane voltage recording (black) and voltage imaging data (red). Shown are data from red-highlighted cell in ***a***. (***c***) Voltage imaging timeseries data from other cells in the same field-of-view as cell in ***a*** and ***b***, including an adjacent cell (purple), which did not show changes in fluorescence when patched cell was injected with current. This indicated no spatial cross-talk of signal. (***d***) Schematic example of showing how response signal-to-noise ratios (SNR) were calculated for trial-averaged fluorescence PSP traces using the mean response amplitude (m), the mean baseline signal (μ), and the standard deviation of bootstrap-sampled baseline means (σ). (***e***) Line plot highlighting the linear relationship between response SNR and the square-root of the number of photostimulation trials averaged. A "detectable" response had an SNR of greater than 2.576. Error bars show SEM, and shaded areas around the fit line show 95% CI of fit.

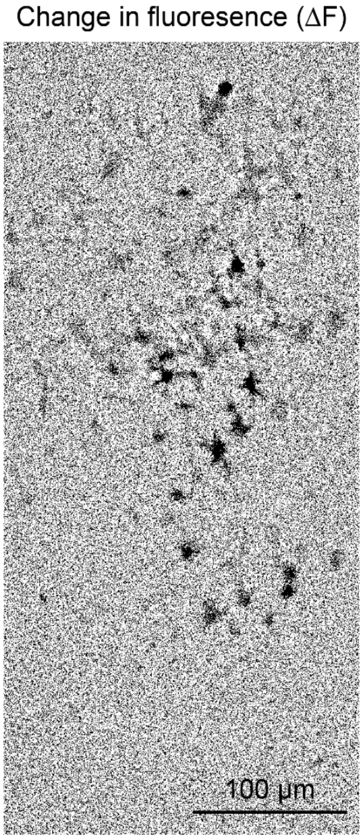


*Figure S2 – Spatial map of fluorescence change following presynaptic photostimulation.* Trial-averaged image of an example MOSAIX field-of-view shows the change in fluorescence (ΔF) of Voltron2-JF585 caused by photoactivation of presynaptic terminals. Pixel intensities have been summed across the 250 ms period following the stimulus. Darker pixels represent larger amplitude PSP responses.

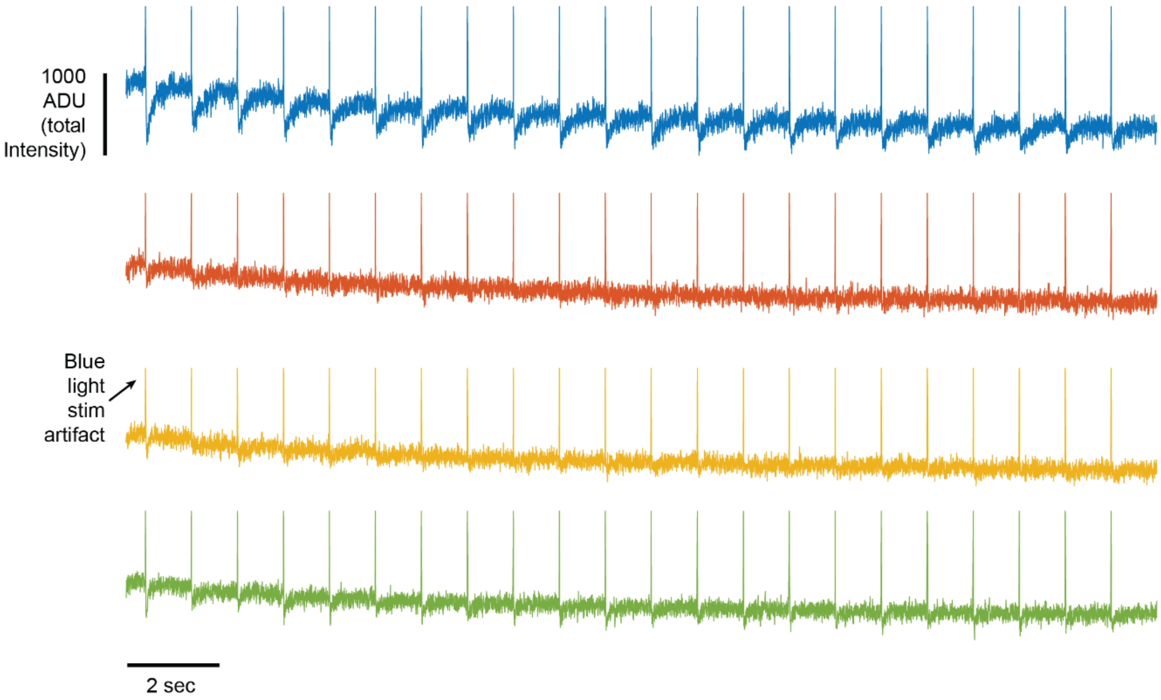


*<u>Figure S3 – Raw fluorescence time series extracted from imaged cells in a MOSAIX experiment.</u>* Raw Voltron2-JF585 signal from four individual cells reveals postsynaptic responses triggered by photoactivation of presynaptic terminals. The blue light stimulus artifact can be measured as a transient increase in fluorescence on a millisecond timescale (upward spikes), while PSPs cause decreases in fluorescence intensity. All data are shown as absolute intensity values.

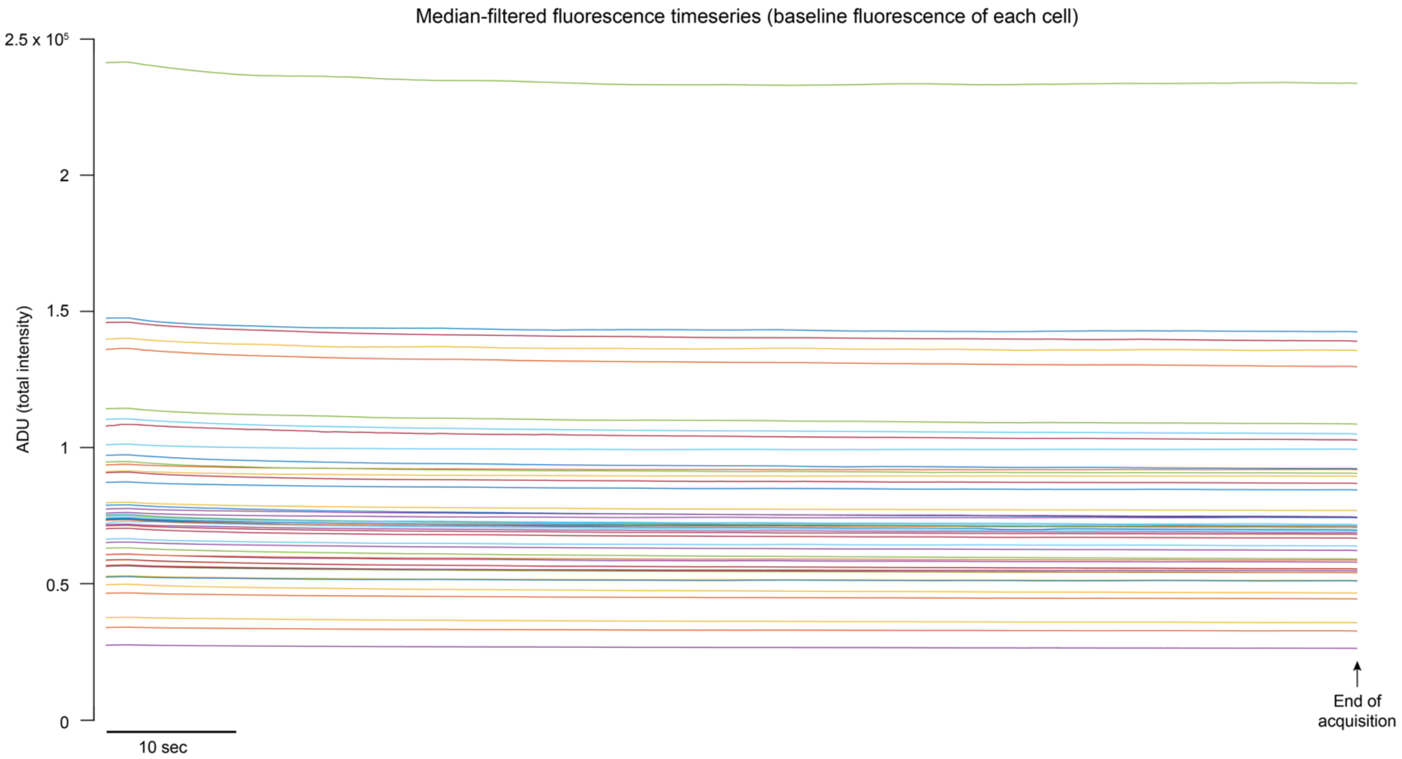


*<u>Figure S4 – Photobleaching during GEVI imaging was minimal.</u>* Traces show the baseline fluorescent signal from individual cells over the length of a MOSAIX data acquisition (each cell shown in a different color). Baseline signal was calculated by filtering a MOSAIX timeseries with a rolling median filter using a 1000 timepoint window. Note that even at the end of an acquisition,

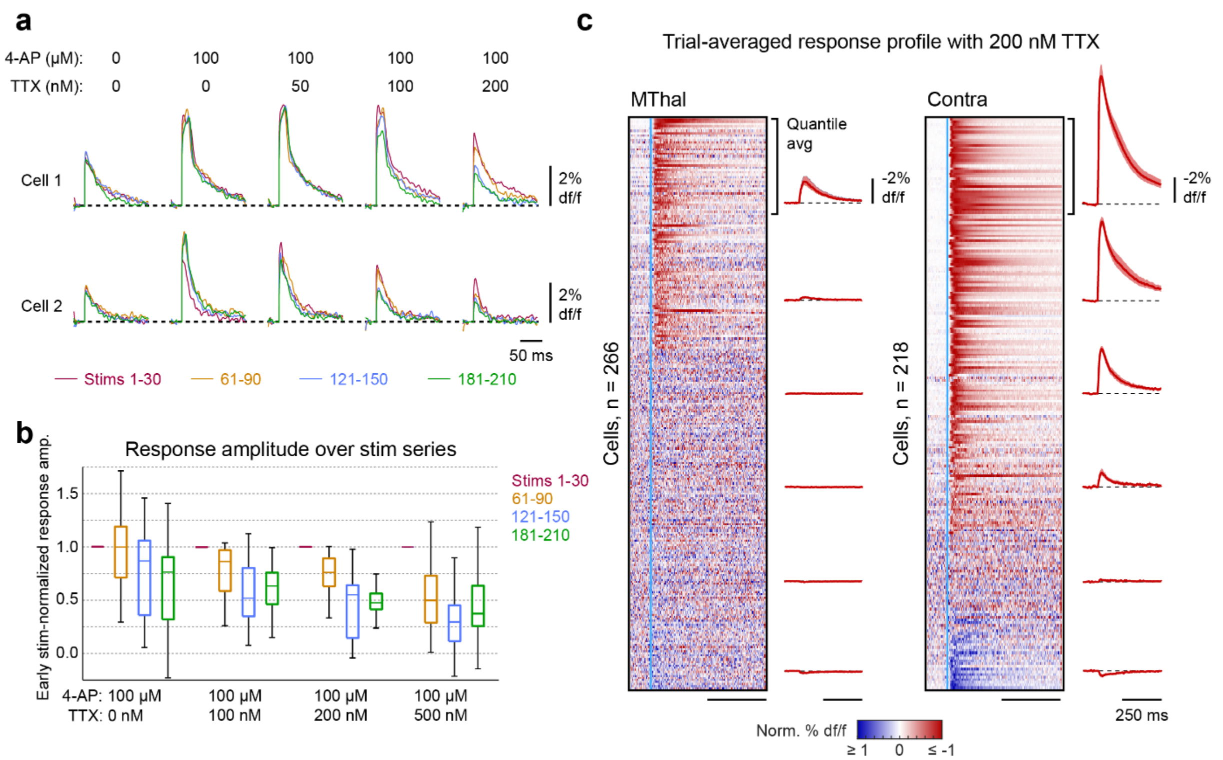


*Figure S5 – Pharmacological agents affect stationarity of repetitive PSPs.* (***a***) PSP timeseries from fluorescence data of two Voltron2-positive cells across pharmacological conditions using same photoactivation power across all conditions. Addition of 100 μM 4-AP increases depolarization in response to presynaptic photoactivation. With increasing concentrations of TTX, cells show attenuation of PSP response amplitudes over the course of the stimulus series. (***b***) Box and whisker plots show extent of non-stationarity across pharmacological conditions, with greatest response attenuation observed with the addition of 500 nM TTX (n = 19 cells). (***c***) Heatmaps showing PSP timeseries across individual cells (rows) imaged in the same experiment, ordered by response amplitude. The traces showing the average PSP within quantile groups are shown alongside (shaded regions show 95% CI). In MThal-to-motor cortex optical connectivity mapping experiments (left), application of 200 nM TTX is sufficient to prevent action potentials and reduce the likelihood of polysynaptic input, whereas in Contra-to-motor cortex optical connectivity mapping (right), 200 nM TTX with the same stimulus intensity results in large-amplitude responses as well as an increase in inhibitory responses, indicating potential polysynaptic signaling.

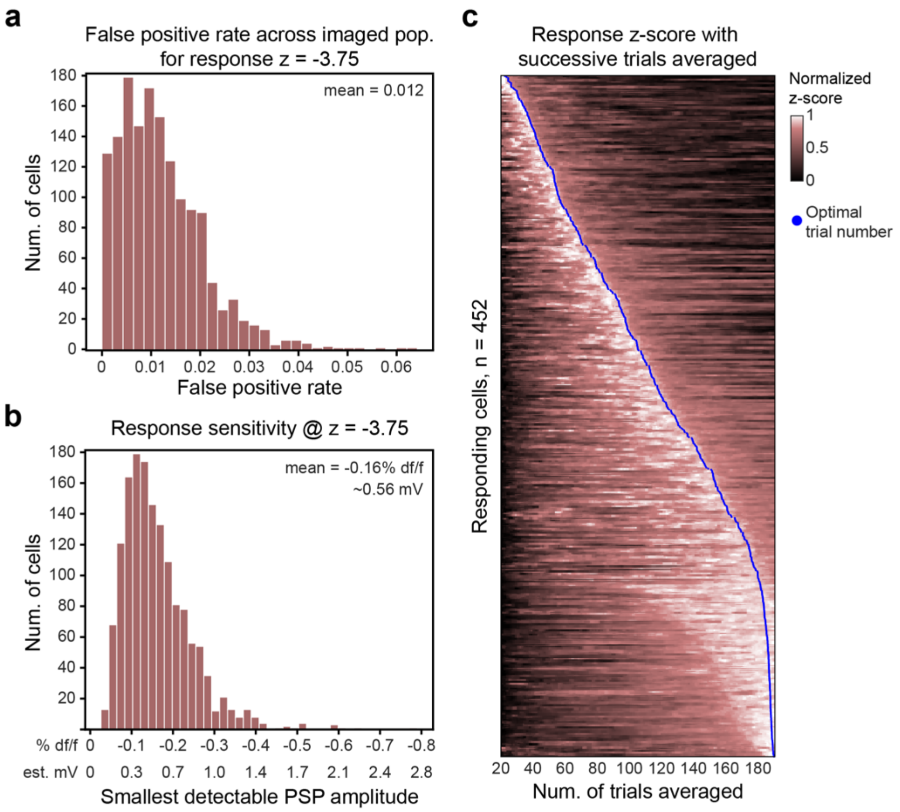


<u>*Figure S6 – Determination of the optimal number of photoactivation trials to average per cell.*</u> (***a***) Histogram showing the false positive rate (FPR) for detection of a postsynaptic response with a z-threshold of -3.75 across all cells imaged in MThal and contra motor cortex optical connectivity mapping experiments. Across all cells, the FPR was below 0.07, with a mean FPR of 0.012. (***b***) Histogram showing the smallest PSP amplitude detectable with the detection z-threshold of -3.75 across all cells imaged in optical connectivity mapping experiments after averaging the optimal number of photoactivation trials (mean = -0.16% df/f, 0.56 estimated mV; n = 1142 cells). (***c***) Heatmap showing the relative z-score of PSP response amplitudes across "responding" cells (rows) with increasing number of trials averaged. The optimal number of trials to average for each cell is highlighted by a blue dot.

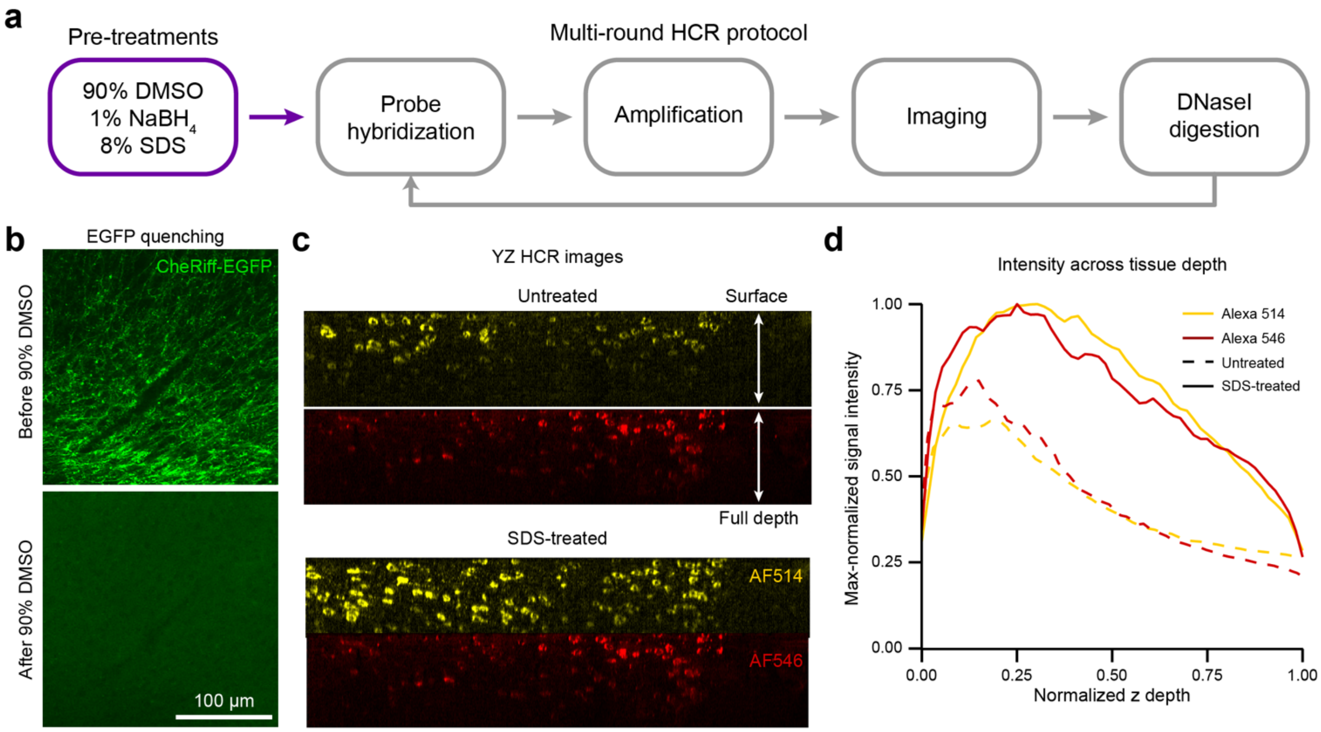


<u>*Figure S7 – The effect of pre-treatment washes on HCR mFISH in 300 µm-thick tissue samples.*</u> (***a***) Outline of custom multi-round HCR mFISH protocol, with pre-treatment steps highlighted in purple. (***b***) Confocal images showing EGFP fluorescence from CheRiff-EGFP fusion before sample incubation in 90% DMSO (top), and absence of EGFP signal following 90% DMSO treatment (bottom). (***c***) Representative orthogonal YZ images through a full-volume confocal z-stack of a 300 µm-thick cortical section. HCR mFISH signal in the AlexaFluor 514 (AF514; yellow) and AF546 (red) channels are shown. Untreated (top) and SDS-treated (bottom) samples are shown. (***d***) Line plots showing quantification of signal intensity across tissue z-depths for AF514 (yellow) and AF546 (red) channels, in both untreated (dotted line) and SDS-treated (solid line) samples.

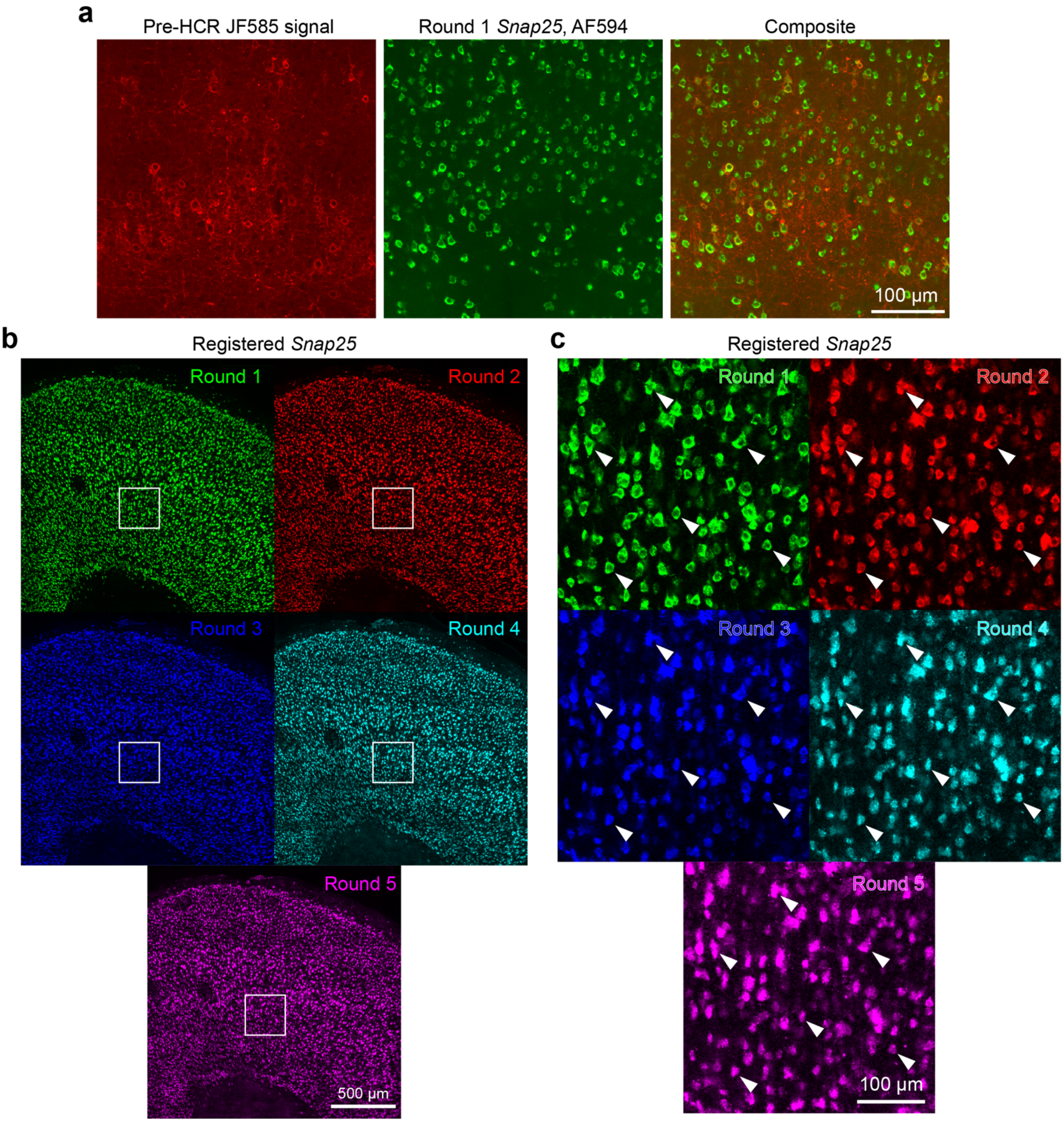


<u>*Figure S8 – Round-to-round registration of mFISH image data using Snap25 signal.*</u> (***a***) HCR pre-treatments extinguish JF585 signal. Images from a registered field-of-view show JF585 signal from pre-HCR confocal imaging (left) with round 1 *Snap25* signal imaged in the AlexaFluor 594 channel (center). Composite (right) shows no signal co-localization. (***b***) Z-projected images of 3 confocal planes, showing registered *Snap25* signal across 5 rounds of HCR mFISH. (***c***) Images from insets shown in ***a***, highlighting individual cells (white arrows) in registered images.

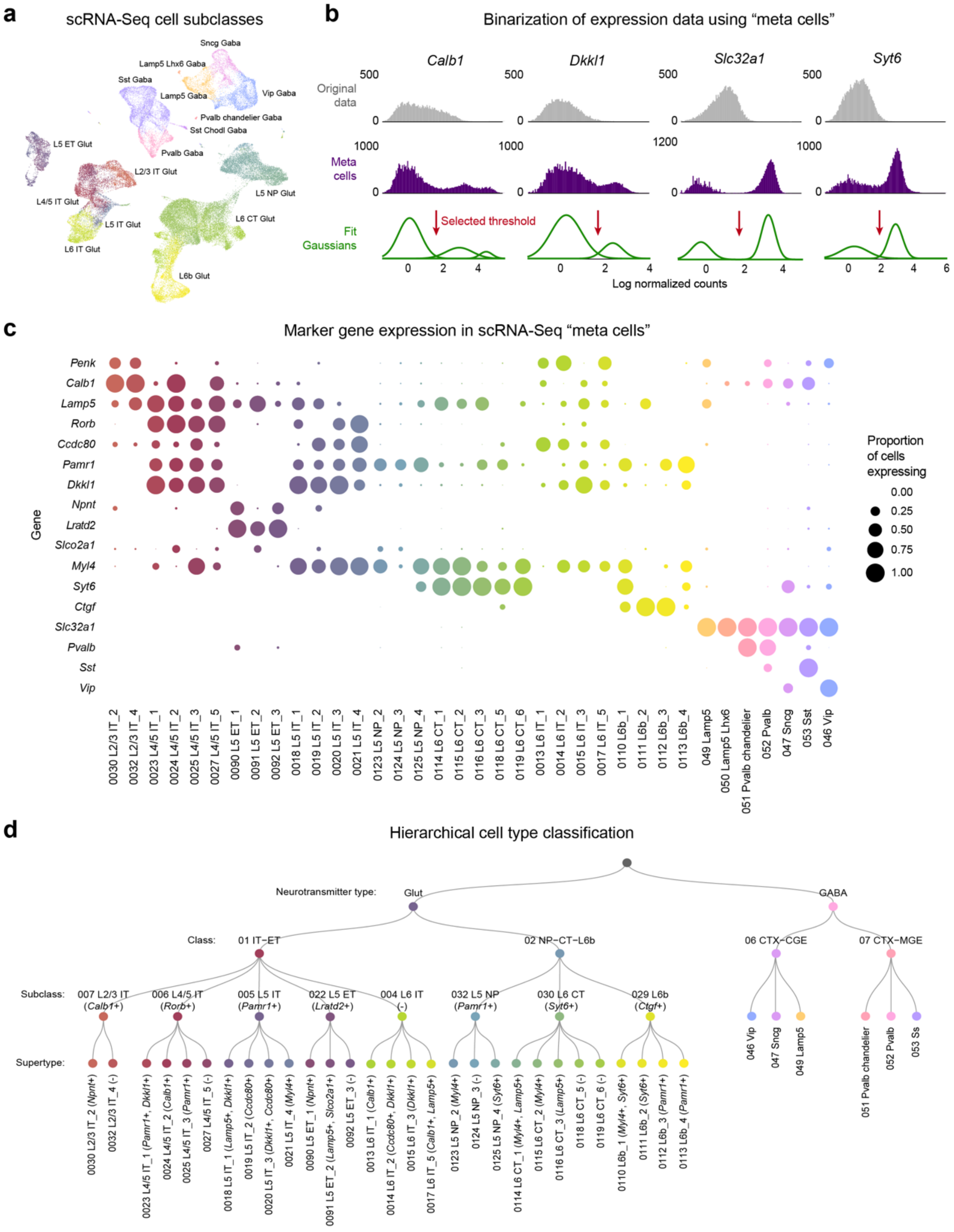


*Figure S9 – Binarization of scRNA-Seq marker gene expression data.* (***a***) UMAP plot generated from ABC Atlas scRNA-Seq data, colored by cell subclass. (***b***) Example histograms for 4 marker genes showing original natural-log normalized counts distributions from ABC Atlas scRNA-Seq data (top, grey), log-norm counts distributions resulting from summing counts from 10 nearest-neighbor cells to create “meta cells” (middle, purple), and the gaussians fit to the meta cell distributions using gaussian mixture models (bottom, green). Gene expression binarization threshold is highlighted with a red arrow, indicating that cells with counts below or above the threshold were considered negative or positive for that gene, respectively. (***c***) Dot plot showing the proportion of excitatory cell supertypes and inhibitory cell subclasses expressing 17 marker genes. Only cell types found in ABC MERFISH data included. Dot size shows proportion of cells of each type expressing each gene. (***d***) Hierarchical tree of cell type classification derived from ABC transcriptomic data. Using our classifier, cells were classified along the hierarchy, first differentiating “neurotransmitter type”, followed by “class”, “subclass”, and “supertype”. Inhibitory cells were only classified down to the level of subclass. Excitatory supertypes are additionally annotated for marker genes that best distinguish them from their “sibling” cell types.

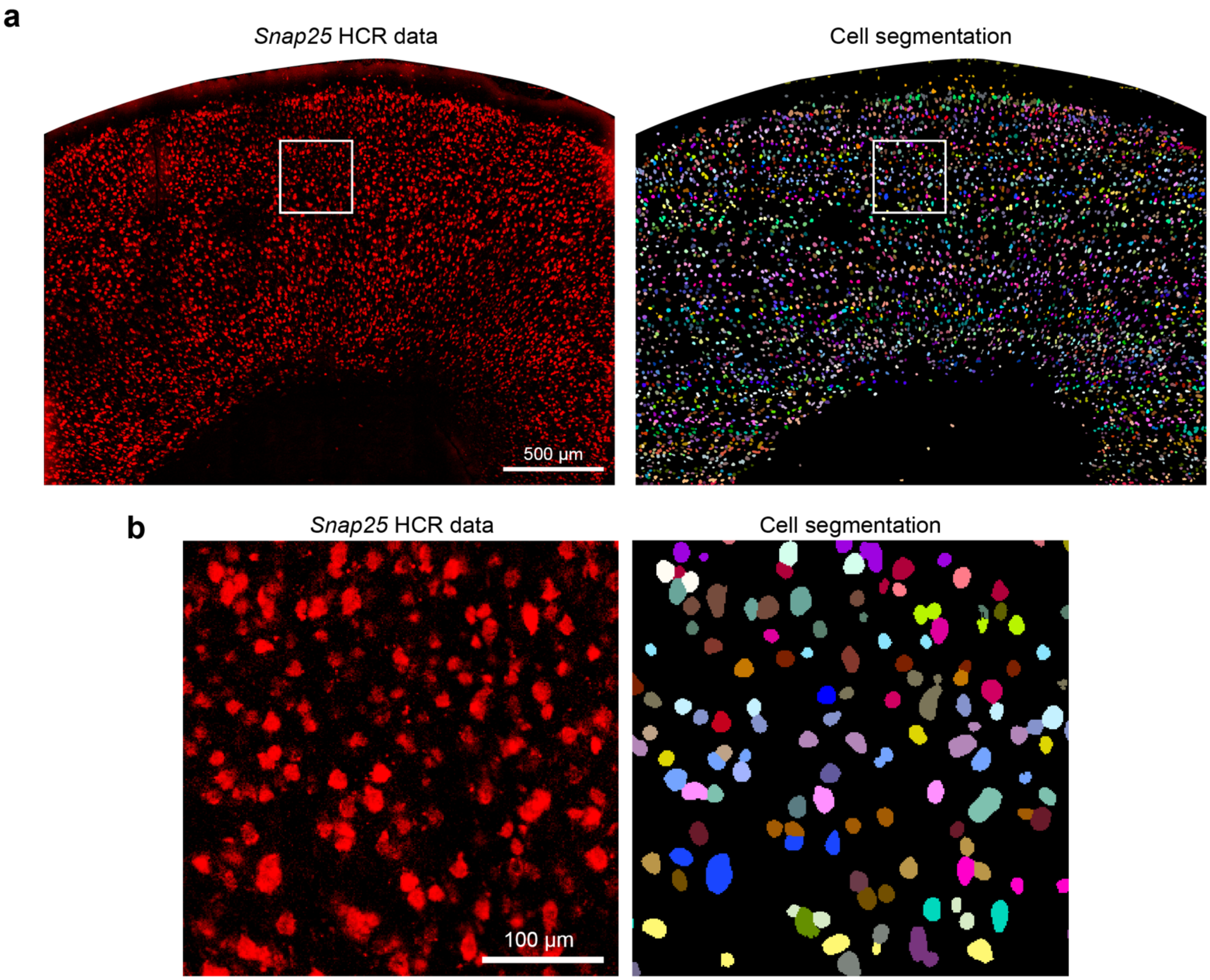


*Figure S10 – Cell segmentation using Snap25 signal to generate cell ROIs.* (***a***) Confocal image of *Snap25* signal in a tissue section of dorsal cortex (left) alongside Cellpose3 segmentation of *Snap25* signal into individual cell ROIs (right). (***b***) Insets of images from ***a***. Cellpose3 allows segmentation of even densely labeled cells.

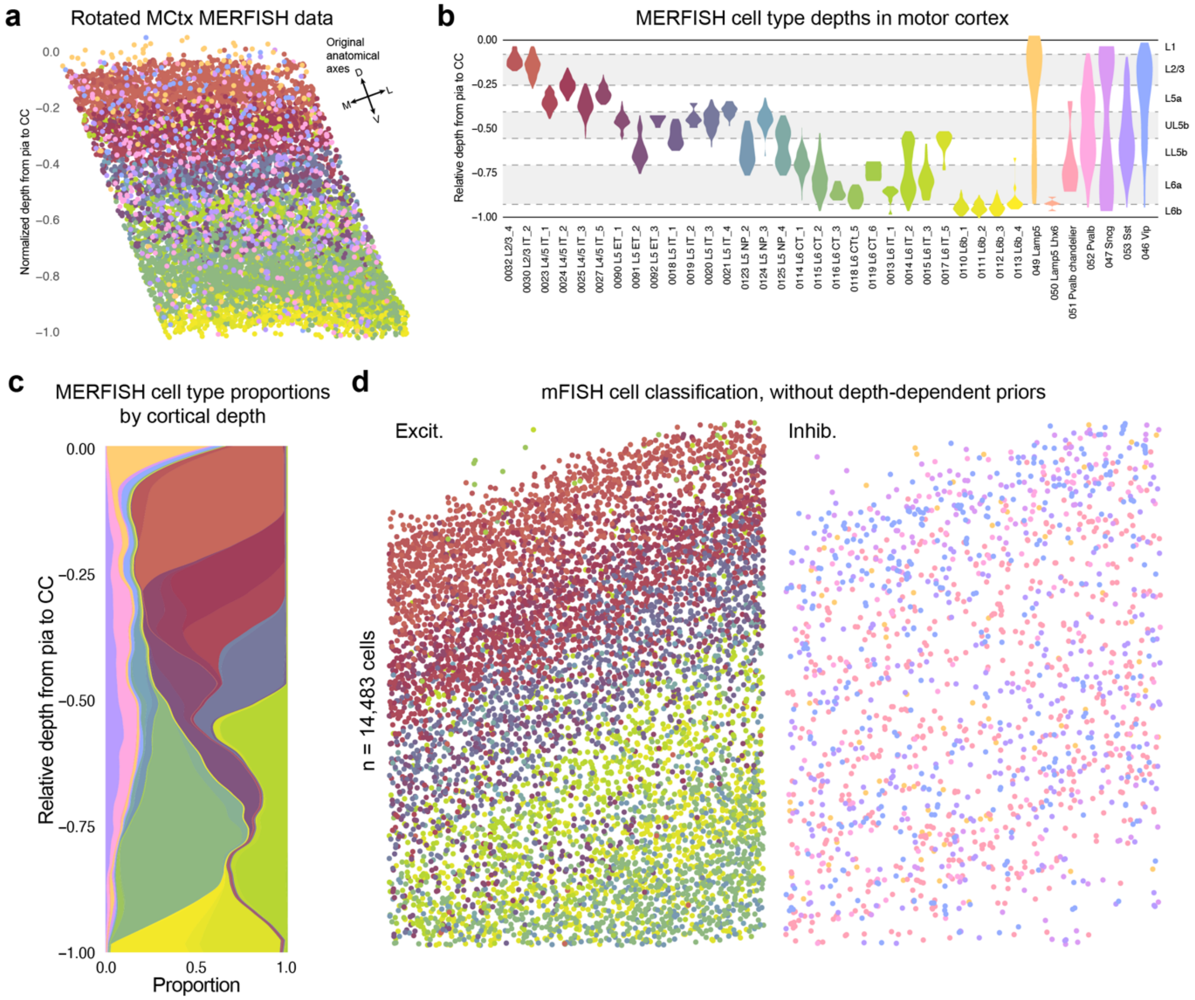


*Figure S11 – ABC MERFISH and scRNA-Seq data provide classification priors.* (***a***) Scatterplot showing arrangement of cells (dots) in ABC MERFISH data from the motor cortex, colored by cell type. Data have been rotated to provide a vertical axis to laminar arrangement of cell types. (***b***) Violin plots showing laminar arrangement of cells in ABC MERFISH data, grouped by excitatory supertypes and inhibitory subclasses used for classification of cells in our HCR mFISH data. Distributions show cells between the $1^{st}$ and $99^{th}$ percentiles of depth. (***c***) Relative cell type proportions across the full depth of motor cortex, determined from ABC MERFISH data, with a baseline probability of 0.2% applied to all cell types. At each depth, these proportions provided cell type prior probabilities for our Bayesian classifier. (***d***) Scatterplots showing motor cortex cells (dots) imaged across 50 µm of z-depth in multi-round mFISH and classified into excitatory (left) or inhibitory (right) cell types based on gene expression patterns (n = 14,483 cells). Cell types were determined using a classifier that did not include depth-dependent cell type priors, in contrast to data shown in ***Fig. 3b***.

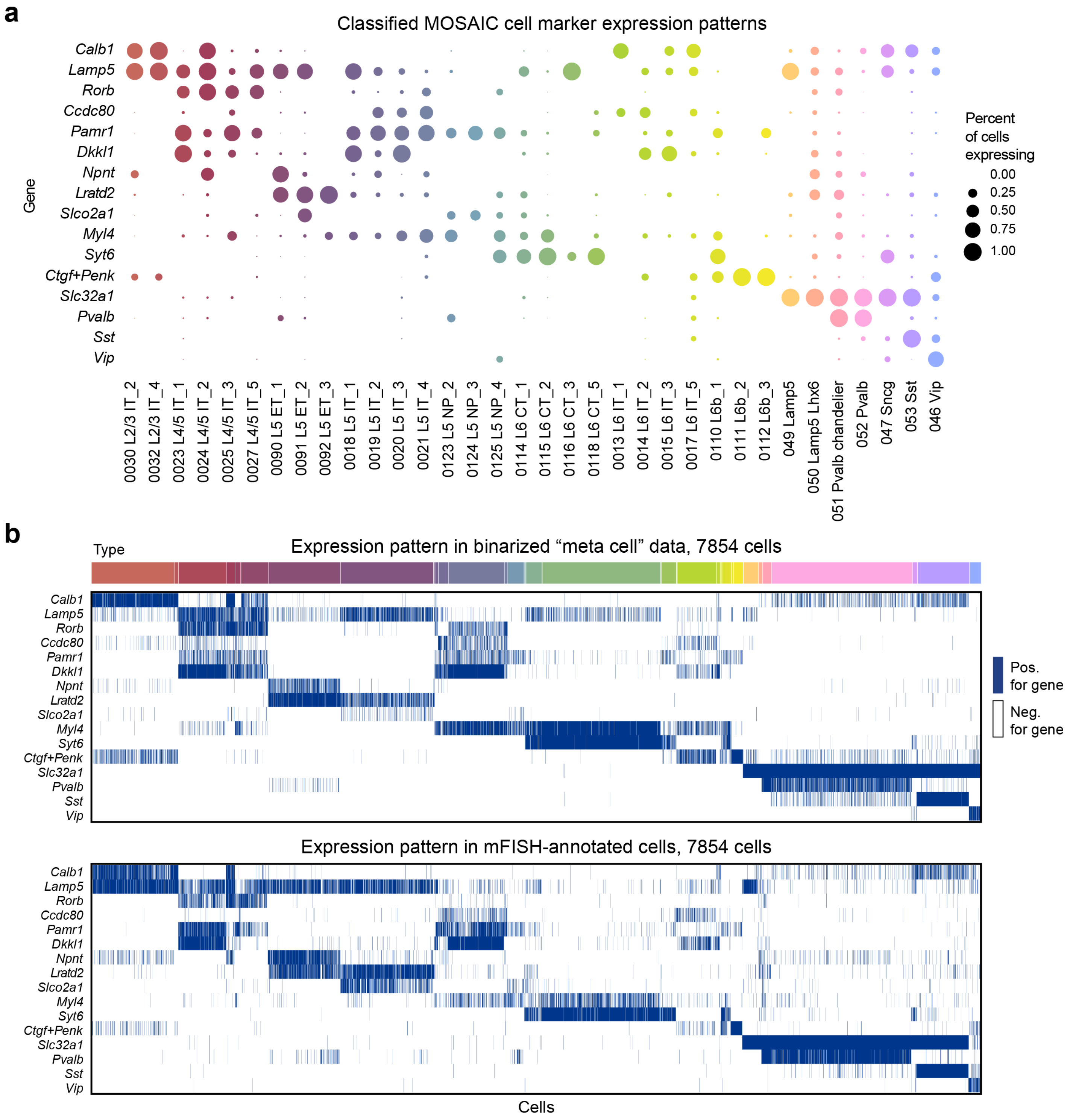


*Figure S12 – Gene expression patterns following classification of MOSAIX data.* (***a***) Dot plot shows marker gene expression patterns of our classified MOSAIX cells. Dot size shows proportion of cells of each type expressing each gene. Only cell types with more than one cell are shown. (***b***) Heatmaps showing the binary expression pattern of marker genes across cells from scRNA-Seq “meta cells” (top) and from annotated MOSAIX image data (bottom). Cells in the “meta cell” dataset were randomly selected to match the number of cells per type found in the MOSAIX data (n = 7,854 cells). Blue indicates where cells were “positive” for a gene, with white indicating where cells were “negative” for a gene.

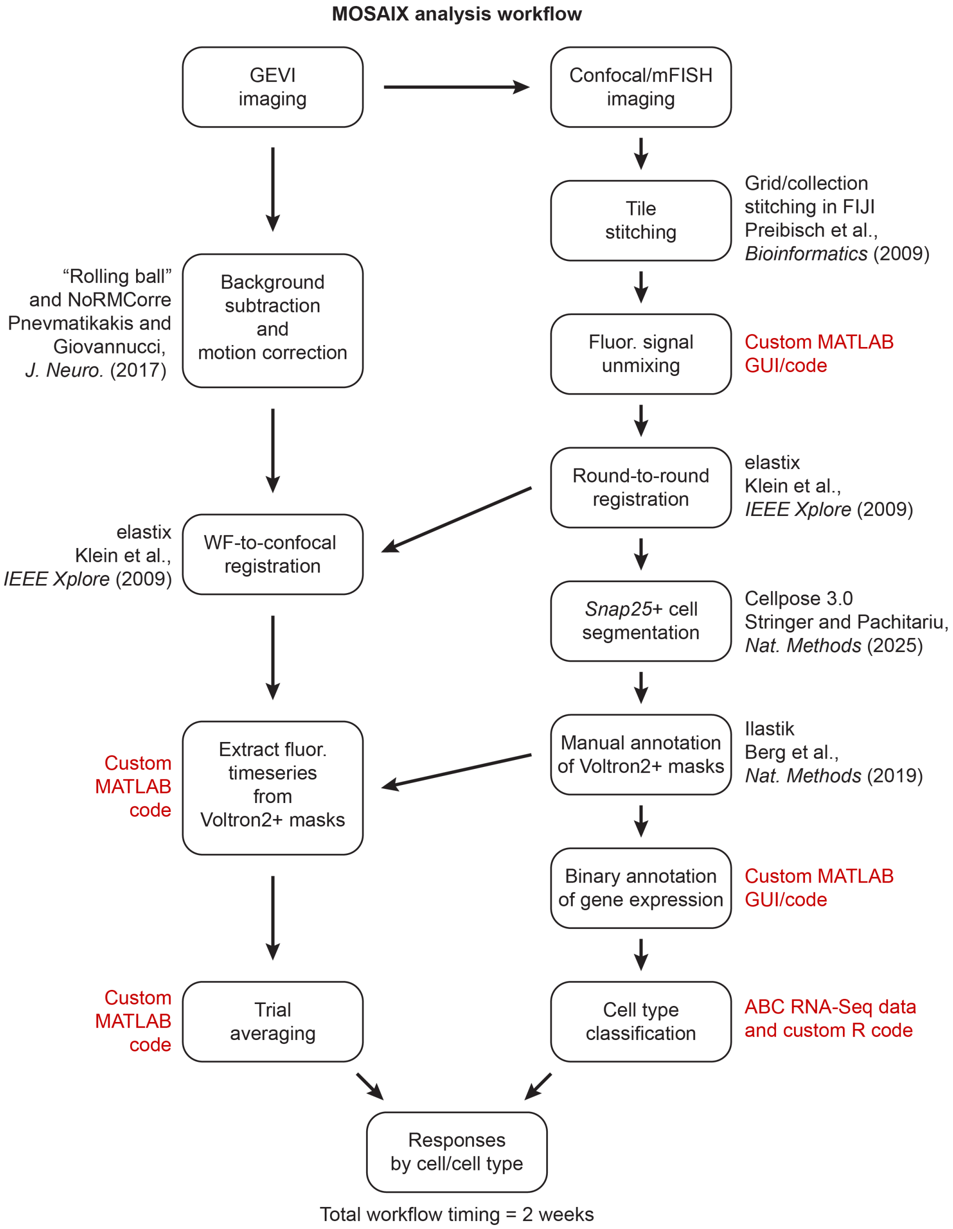


*Figure S13 – MOSAIX analysis workflow.* Outline of the analysis pipeline used for analyzing GEVI imaging and mFISH imaging data in MOSAIX experiments. Steps using publicly available tools have included citations for the original reference to the tools. Steps using custom code, guis, or algorithms have been highlighted in red and are shared as part of this publication.

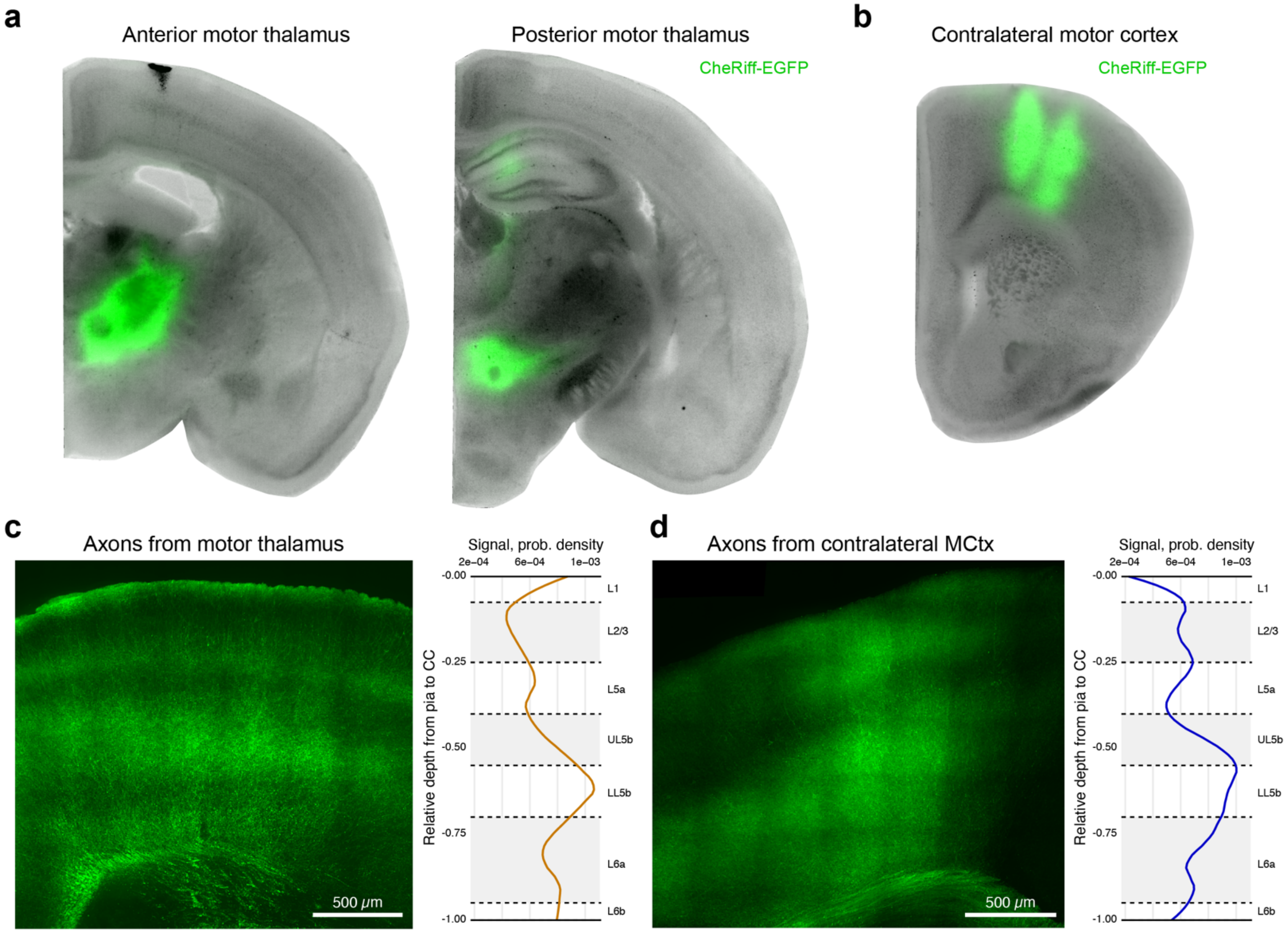


*Figure S14 – Injection site histology for AAV9-hSyn-CheRiff-EGFP delivery to MThal and contra motor cortex.* (***a***) Representative images showing coronal hemisections with AAV9-hSyn-CheRiff-EGFP injections into anterior (left) and posterior (right) MThal. CheRiff-EGFP signal (green) is shown against tissue autofluorescence (grey). Nuclei showing expression include VAL and medial VM. (***b***) Representative image showing injections into two medial-lateral coordinates in contra motor cortex. Injections were additionally targeted to two cortical depths to achieve expression across cortical layers. (***c, d***) Representative image of CheRiff-EGFP+ MThal axons (***c***) and contra motor cortex axons (***d***) in the ipsilateral motor cortex (left). EGFP signal density plots across cortical depths are shown on the right of each image.

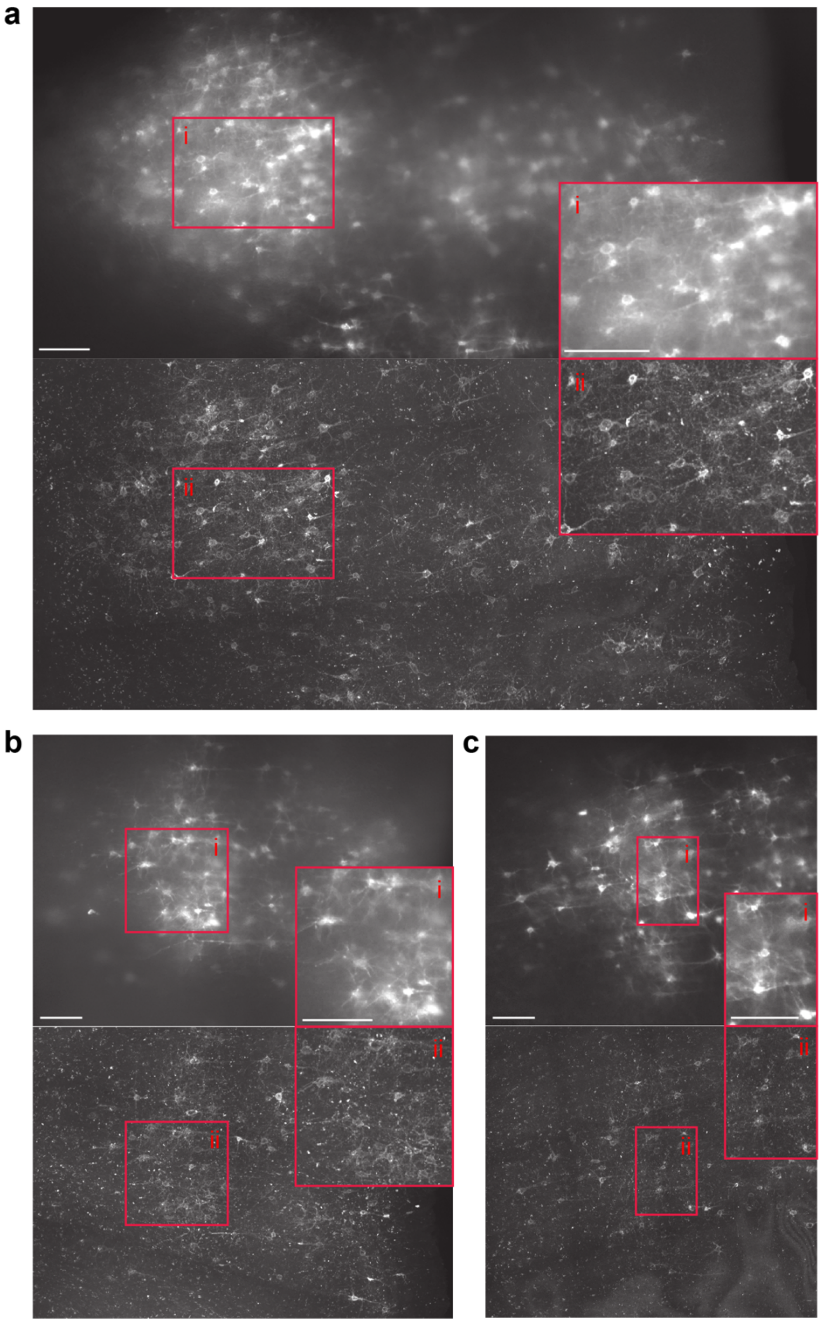


*Figure S15 – Registration of Voltron2 image data in MOSAIX.* (***a-c***) Representative images from separate MOSAIX experiments show widefield Voltron2-JF585 signal acquired during OCM (top) and registered post-hoc Voltron2-JF585 confocal image used for mFISH histology (bottom). Insets of the same region of each image are shown in red boxes. Scale bars show 100 µm.

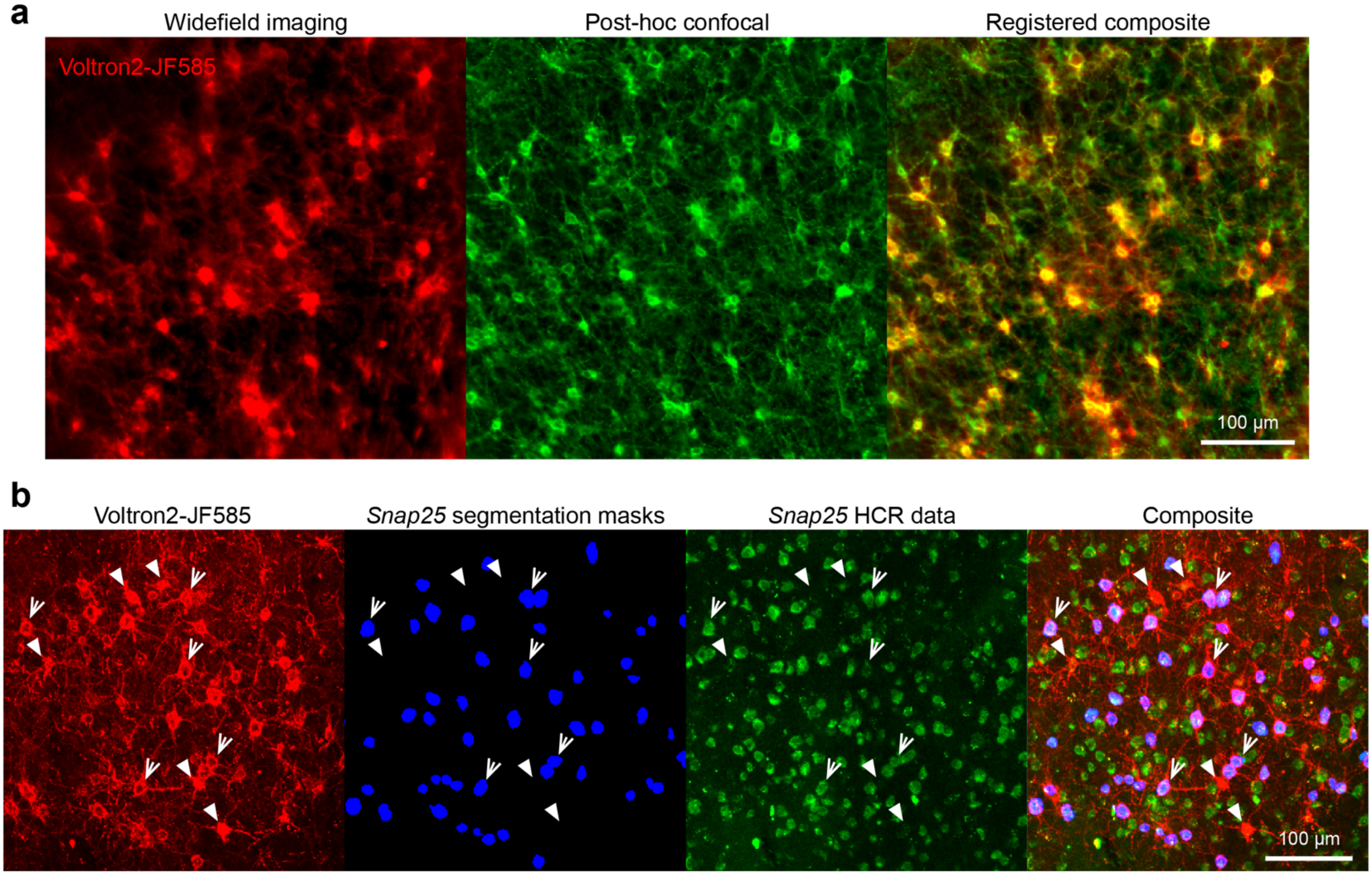


<u>*Figure S16 – Segmentation and registration of Voltron2 image data in MOSAIX.*</u> (***a***) Representative images showing widefield Voltron2-JF585 signal acquired during OCM (red), registered post-hoc Voltron2-JF585 confocal image used for mFISH histology (green), and image composite. (***b***) Image array showing a confocal histology image of Voltron2-expressing cells imaged in an OCM experiment (red), Snap25-based segmentation ROI masks annotated as Voltron2+ (blue), Snap25 mFISH data used for cell segmentation (green), and composite image. Solid arrows highlight Voltron2+ cells that did not show Snap25 signal and corresponding Snap25 segmentation masks, and dashed arrows show Voltron2+ cells that had both.

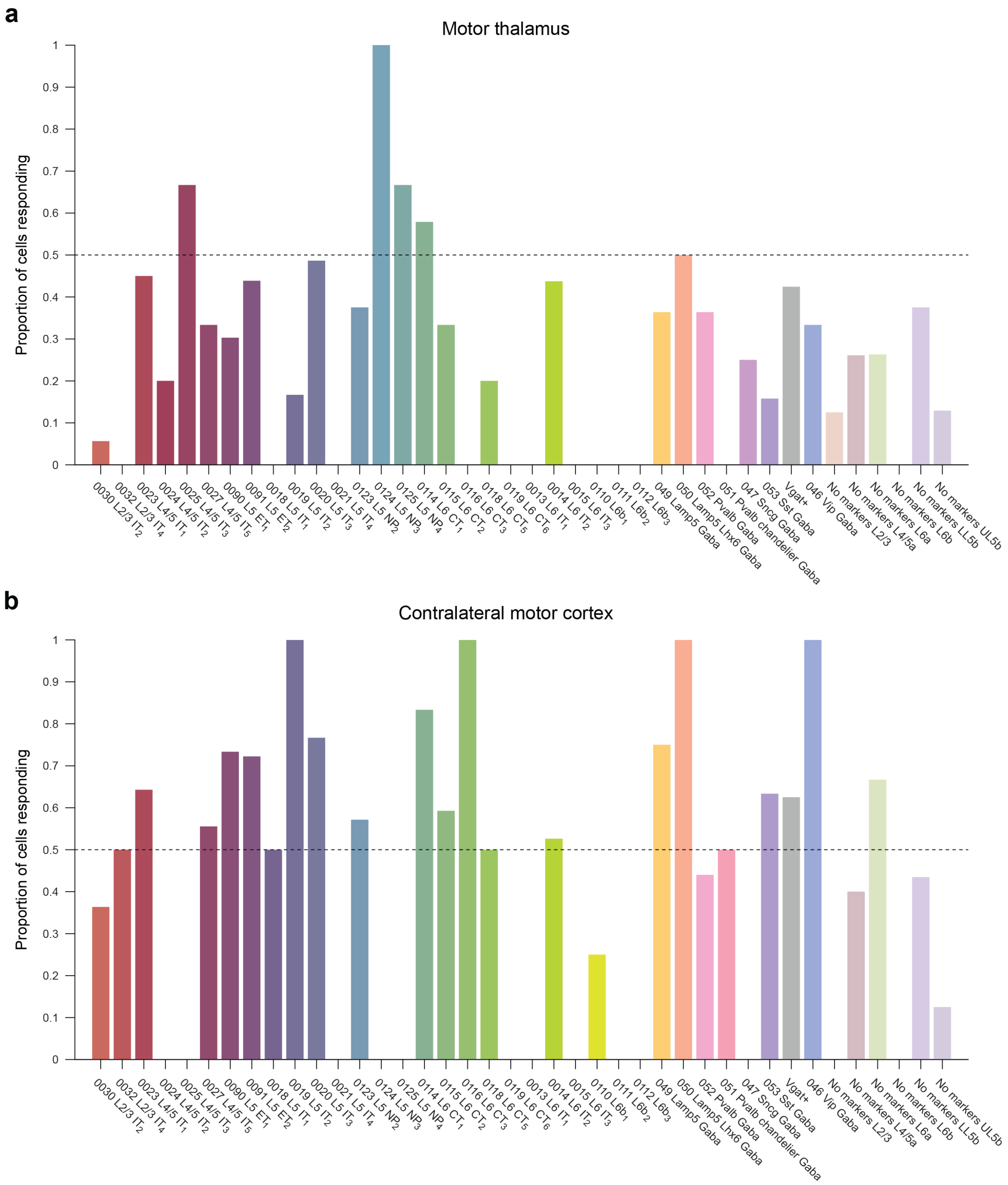


*Figure S17 – Population response rates across cell types.* (***a, b***) Bar charts show the proportion of cells of each type that was classified as "responding" across cell types. Data are shown for motor thalamus (***a***) and contralateral motor cortex (***b***) MOSAIX experiments. Missing bars indicate a cell type that was not sampled in that circuit.

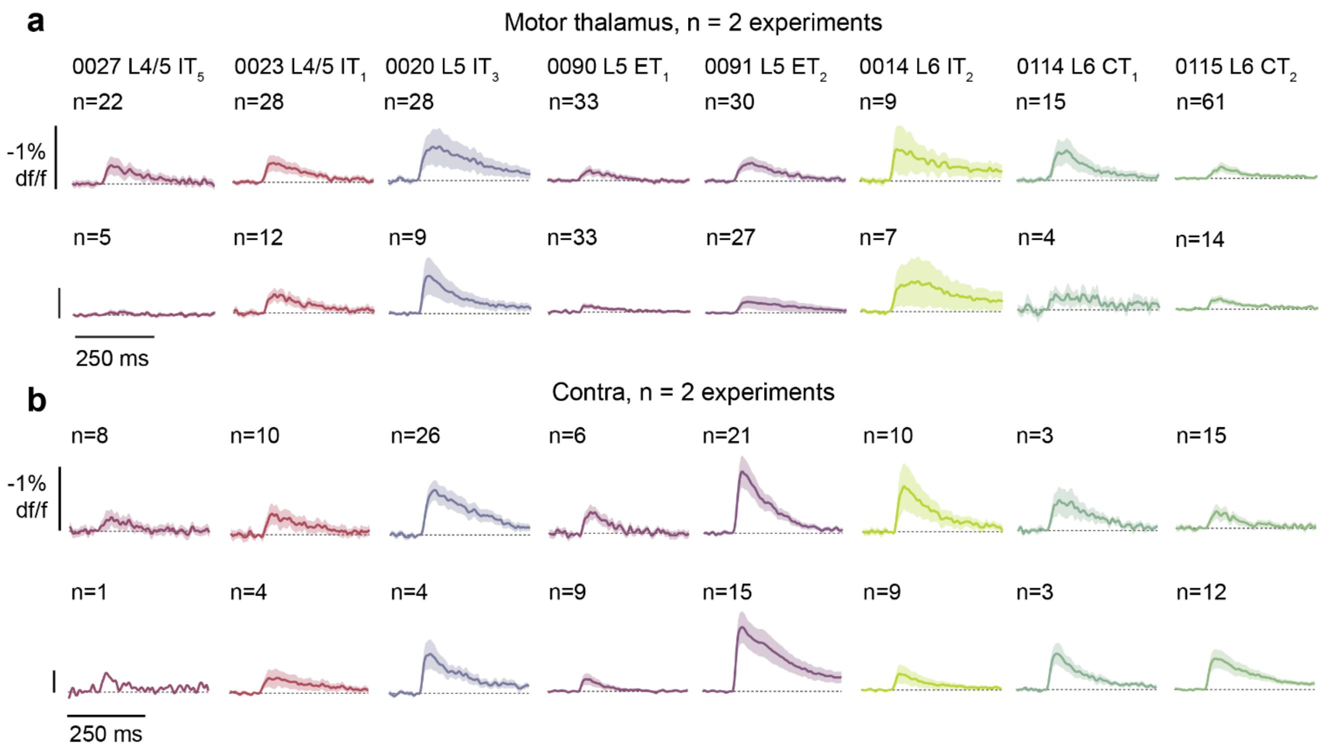


*Figure S18 – Reproducibility across individual MOSAIX experiments.* (***a, b***) Mean PSP profiles (averaged across both "responding" and "non-responding" cells) resulting from activation of MThal inputs (***a***) and Contra inputs (***b***). For each circuit, data from two separate is animals/experiments is shown in two separate rows. Shaded region shows standard error of the mean.

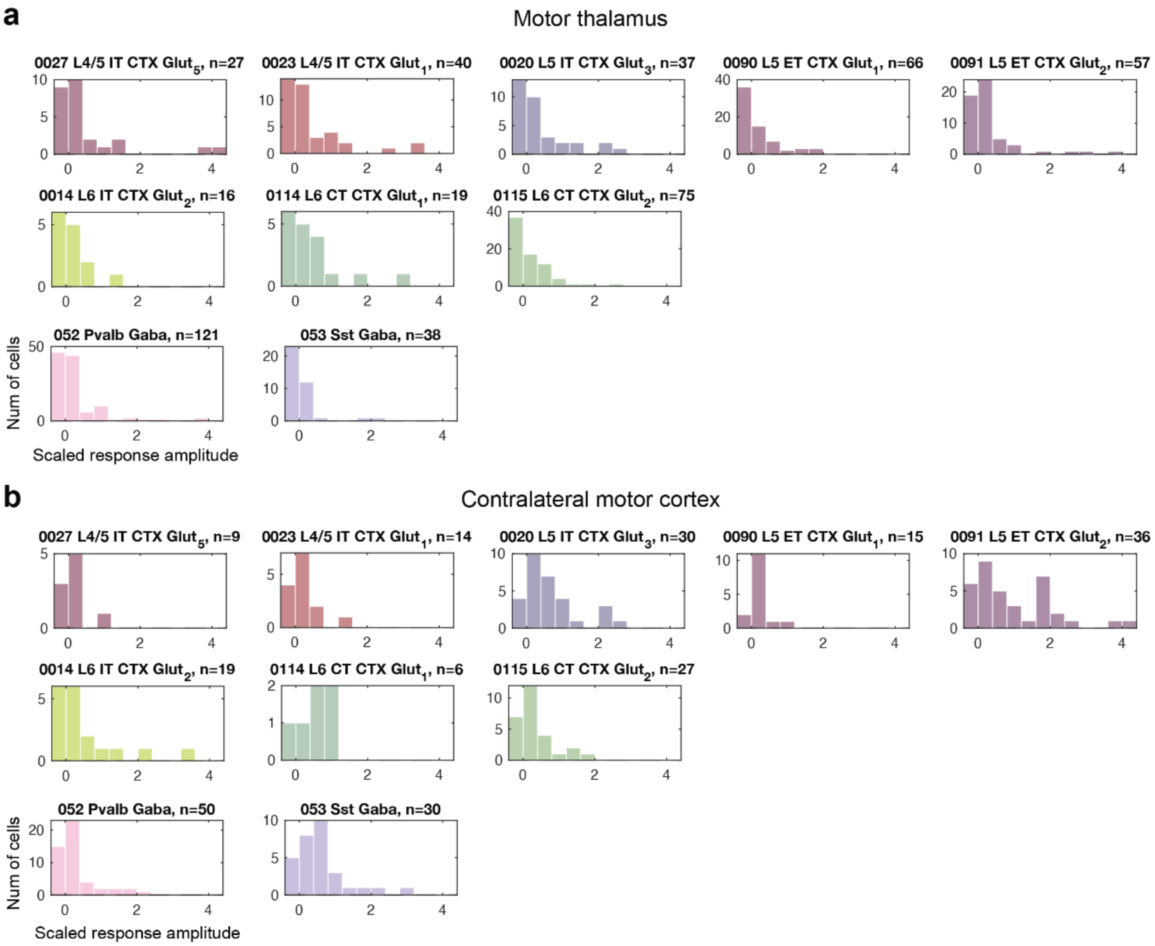


*Figure S19 – Scaled response amplitudes across cell types in MOSAIX data.* (***a, b***) Histograms show response amplitudes across both responding and non-responding cells, scaled by dividing each value in the PSP trace by the mean PSP amplitude of the cell type with the largest response. Data are shown for all MThal (***a***) and contra motor cortex (***b***) MOSAIX experiments across cell types featured in Figure 5 and Figure S18.

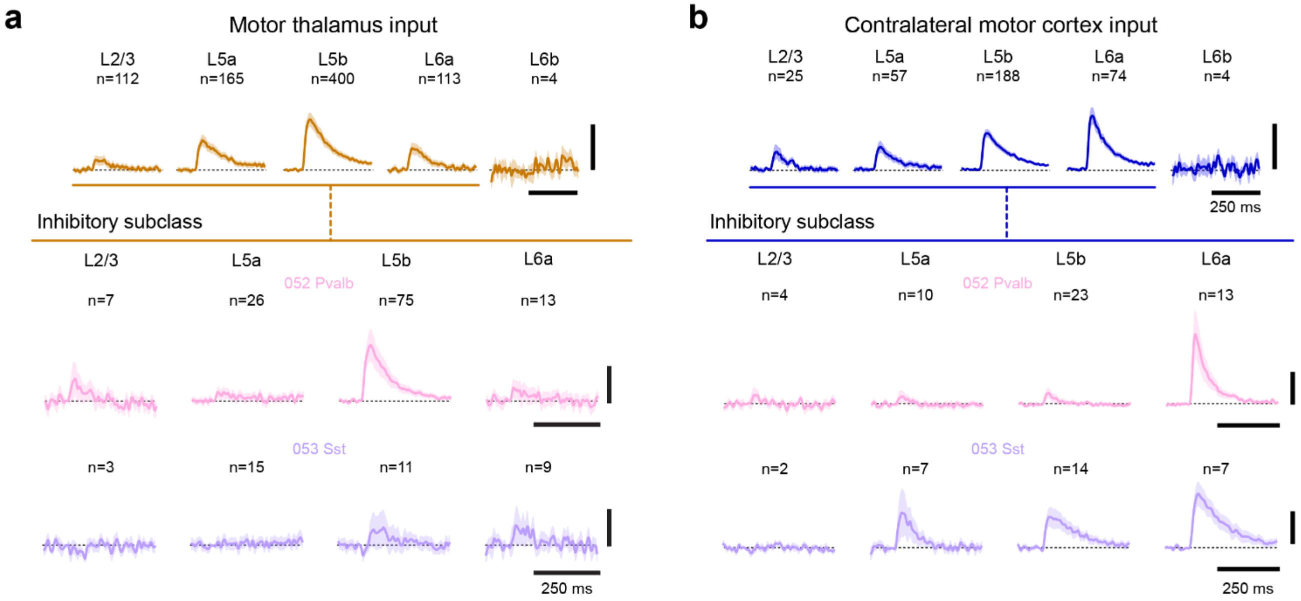


<u>*Figure S20 – MOSAIX data provide connectivity data for inhibitory cell types.*</u> (***a, b***) Scaled motor cortex PSPs resulting from activation of MThal inputs (***a***; orange) and contra motor cortex inputs (***b***; blue), grouped across all cell types by laminar depth (top – same traces shown in main Fig. 5), and across 052 Pvalb and 053 Sst inhibitory subclasses for cells found in L2/3-L6a (bottom). Line shows mean trace, shaded area shows SEM. Vertical scale bars show 1x scale.

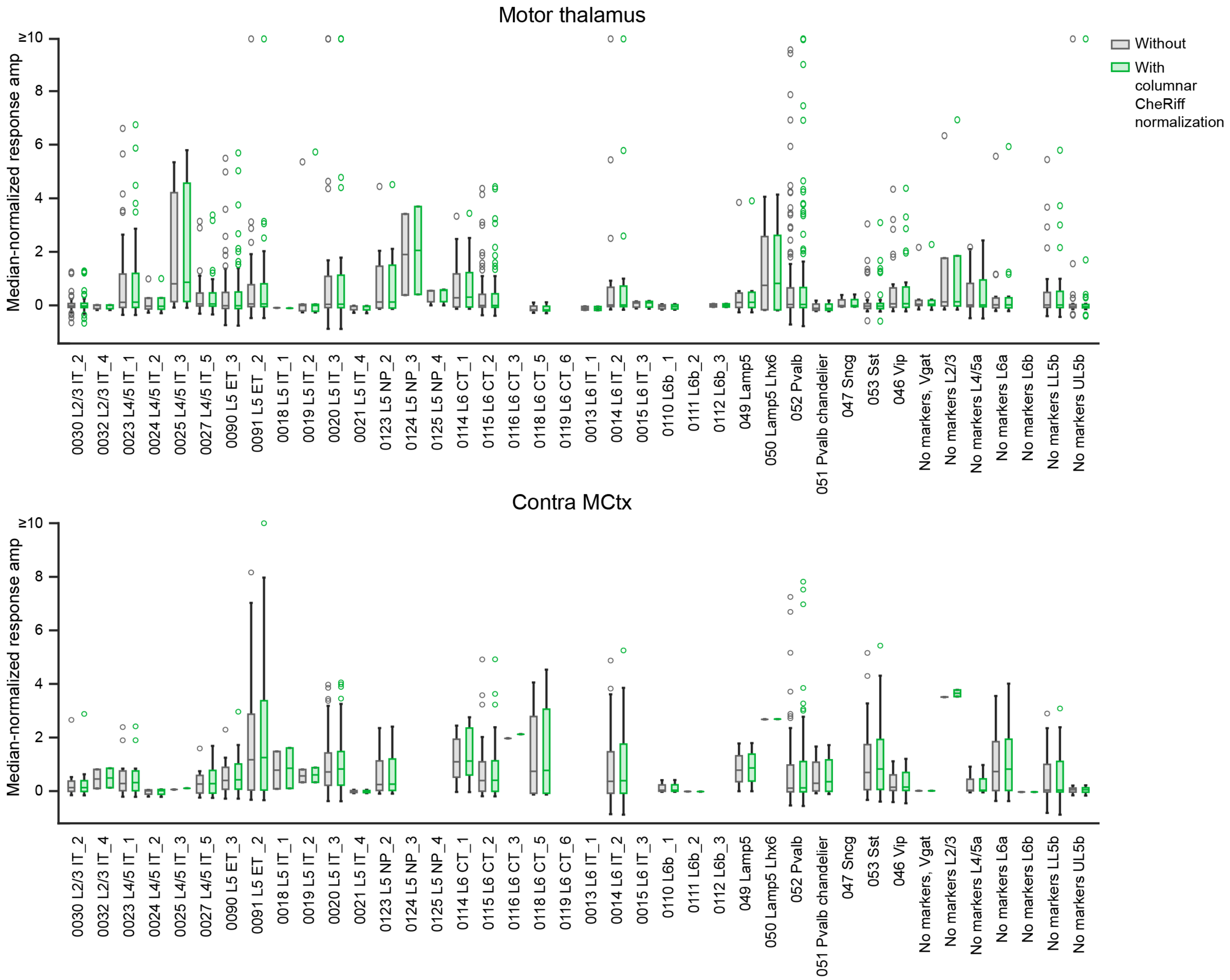


*Figure S21 – Normalized response amplitudes across cell types in MOSAIX data.* Box and whisker plots showing response amplitudes across both responding and non-responding cells, normalized to the median response of responding cells in the same experimental slice (grey), or to both the median response amplitude and the relative CheRiff-EGFP signal found in the cell's cortical column (green). Data are shown for all MThal (top) and contra motor cortex (bottom) MOSAIX experiments across cell types.
